# SysNDD: A Systematic Database for Neurodevelopmental Disorders

**DOI:** 10.64898/2026.09.29.755401

**Authors:** Bernt Popp, Simon Früh, Melek Firat Altay, Hilde Van Esch, Almuth Caliebe, Nuria C. Bramswig, Fabian Hummel, Sopio Gverdtsiteli, Tjitske Kleefstra, Annette Schenck, Zeynep Tümer, Alain Verloes, Christiane Zweier

## Abstract

Neurodevelopmental disorders (NDDs) are clinically and genetically heterogeneous, and evidence linking additional genes to disease evolves rapidly. We developed SysNDD, an open-access, expert-curated resource that represents distinct gene-inheritance-disease associations as entities with standardized confidence, phenotype, and variant annotations. Each entity retains its curation history, so its evidence category can change as evidence accumulates. Currently, SysNDD contains 4,275 entities curated from 4,932 publications. Of these, 3,758 entities involving 3,271 genes describe NDDs; the remainder record non-NDD disorders, nearly all linked to NDD-associated genes, included for completeness. Regarding NDD evidence categories, 1,838 genes are Definitive, 179 Moderate, 1,248 Limited, and 6 Refuted. The entity model captures inheritance-specific disorders, including 220 genes with both autosomal dominant and autosomal recessive NDD associations. Comparison with seven other NDD gene resources showed complementary coverage, and evidence categories largely agreed with those of ClinGen, Gene2Phenotype, and PanelApp for shared gene-disease pairs. NDD genes formed distinct molecular modules, whereas associated disorders formed broad, overlapping phenotype groups that corresponded to those modules only selectively. SysNDD provides a continuously maintained, programmatically accessible framework for gene-disease evidence assessment, inheritance-aware interpretation, and systematic NDD research.

## Introduction

High-throughput sequencing is now considered standard for diagnosing rare genetic diseases. It has been particularly successful in neurodevelopmental disorders (NDDs), where trio-based exome or genome sequencing identifies a causative variant in more than 40% of cases^1^. Large-scale *de novo* and rare-variant studies continue to expand the candidate-gene landscape for neurodevelopmental delay and autism^2,3^. Through large consortia and matchmaking platforms, international collaborations have identified many new NDD-associated genes and raised the diagnostic yield^4^. New classes of disease genes also continue to emerge, including non-coding genes such as *RNU4-2*, which encodes the spliceosomal U4 snRNA^5^. As a result, NDDs are among the most clinically and molecularly heterogeneous groups of monogenic disease, with one of the most rapidly evolving genetic landscapes.

This heterogeneity is evident when comparing published numbers of NDD genes before and after the advent of large-scale exome sequencing studies such as the Deciphering Developmental Disorders (DDD) study^1^. Before large-scale sequencing, approximately 300 genes had been reported for NDDs^6,7^. As of September 21, 2026, SysNDD, the resource presented here, records more than 1,800 genes with a Definitive NDD association, a number that continues to grow. SysNDD contains more than 1,200 further genes whose strongest NDD association is so far Limited. Gene-disease associations outnumber genes, because a gene can be linked to multiple diseases, not all of which include developmental delay/intellectual disability (ID). Maintaining current, expert-curated, and complementary resources is therefore essential to inform accurate diagnostics and clinical research^8^.

Existing gene knowledge databases include generalized catalogs of rare disease associations such as Orphanet^9^ and Online Mendelian Inheritance in Man (OMIM)^10^, diagnostic gene panels (e.g., PanelApp^11^), and specialized NDD-focused gene-validity resources such as Gene2Phenotype^12^ and SFARI^13^. However, these curation efforts differ in scope, evidence standards, and interoperability. Standardization initiatives such as the ClinGen framework^14,15^ and the Gene Curation Coalition (GenCC)^16,17^ have helped harmonize gene-disease validity assessment across resources. Against this background, SysNDD provides a disease-centered NDD resource with standardized phenotype and variant annotations, evidence categories broadly comparable to ClinGen/GenCC terminology, collaborative expert curation, and full programmatic access through a public application programming interface (API) that software can query directly.

Here, we present SysNDD (Figure 1), an extensive further development of the previous SysID database^18^ with a new database concept, data model, and infrastructure. We use this resource, structured around gene-inheritance-disease entities, to delineate the current NDD landscape and to ask whether NDDs still resolve into the coherent phenotypic and functional modules previously described for intellectual disability^18^. Four questions guide the analysis: how many NDD genes are known and how the catalog grows; how SysNDD compares with other gene resources; which phenotypes and variant types are recorded; and whether clinically similar disorders involve genes that interact molecularly.

**Figure 1:**
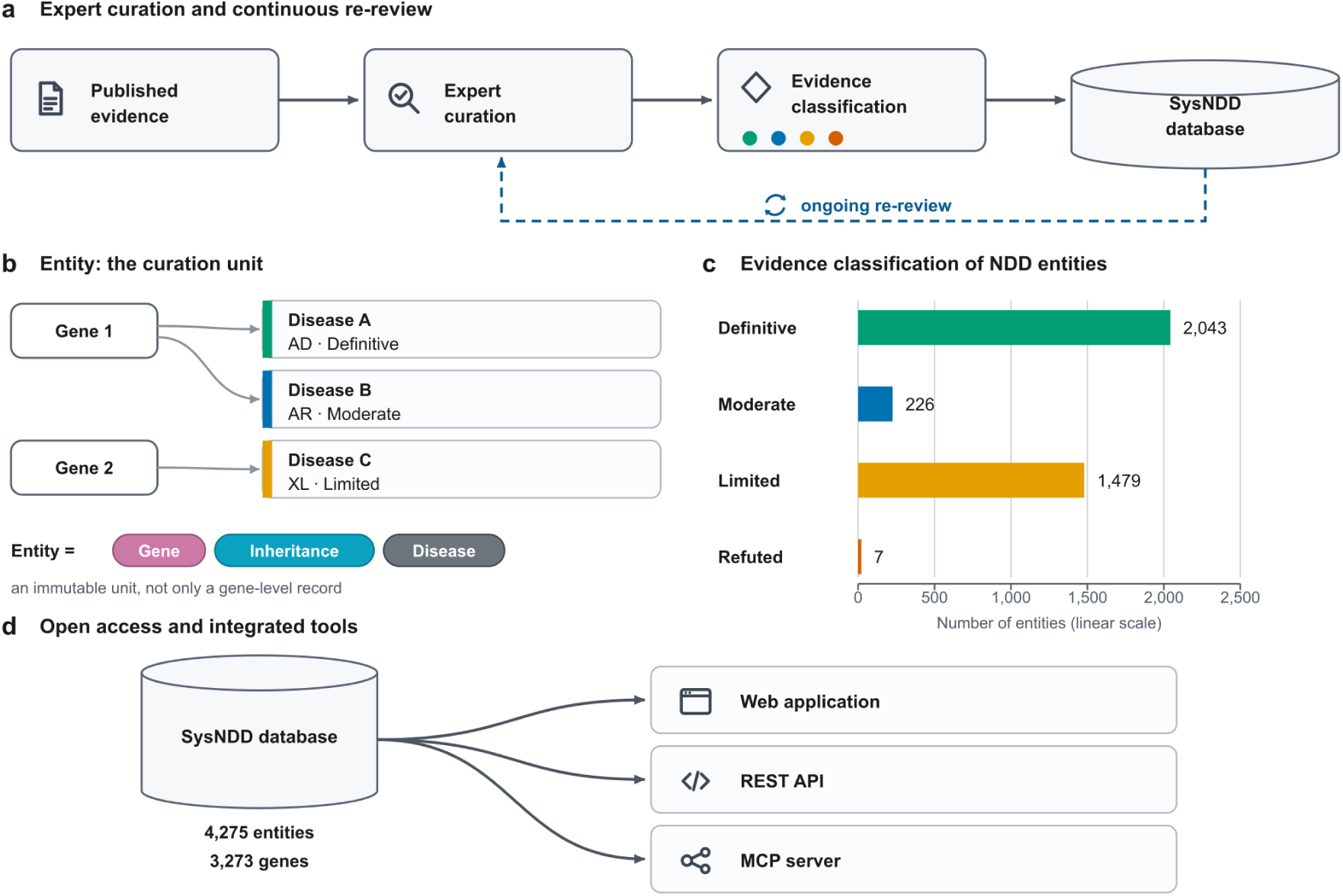
The SysNDD concept, curation approach, and tools. **a**, Curation workflow: evidence from literature is expert-curated (including annotation of phenotypes, variant and inheritance data), assigned to one of four evidence categories (Definitive, Moderate, Limited, or Refuted), and re-reviewed by ERN-ITHACA experts on an ongoing basis; all records are standardized to community ontologies (HGNC, HPO, OMIM, MONDO, and VariO). **b**, The curation unit is the entity, an immutable gene-inheritance-disease triple rather than a gene-level record; one gene may yield several entities, including dual-inheritance (Gene 1) or X-linked (Gene 2) associations. **c**, NDD-associated entities by evidence category (bar length proportional to entity count, linear axis): 2,043 Definitive, 226 Moderate, 1,479 Limited and 7 Refuted; the 3 uncategorized NDD entities are not shown. **d**, The openly licensed database (4,275 entities, 3,273 genes, 4,932 publications; CC BY 4.0 data) is accessible through a web application, a documented REST API, and a read-only Model Context Protocol server. Data: SysNDD API v0.38.1 (accessed September 21, 2026).

## Methods

### The entity data model

SysNDD is based on the “entity” concept (Figure 1b): a fixed triple of one gene, one mode of inheritance and one disease, in which pathogenic variants in that gene cause that disease through that inheritance pattern. Entities use standard identifiers: HUGO Gene Nomenclature Committee (HGNC) for genes^19^, Human Phenotype Ontology (HPO) for inheritance^20^, OMIM for disease names^10^, and Mondo Disease Ontology (MONDO) where no OMIM entry exists^21^. Each entity carries an evidence category (Definitive, Moderate, Limited, or Refuted; Table S1) reflecting the strength of the gene-disease association at the time of curation. Entities for non-NDD diseases linked to an NDD-associated gene are tagged “Not applicable”, since they fall outside the NDD evidence categories, and are kept for completeness. Entities are annotated with 38 HPO phenotype categories, published literature references, variant types and consequences using the Variation Ontology (VariO)^22^, and a textual summary. Variant annotations are assigned by curators from the primary literature. Alongside the curated entities, the web interface displays external reference data for the same gene, namely gnomAD gene constraint metrics^23^ and ClinVar variant records^24^; these data are not imported into the curated entity. Curation history is retained for each entity. Re-review adds new review and status records, and the newly approved status becomes the active one. Entities are never deleted. Associations that no longer meet the criteria are demoted or reclassified as Refuted and remain visible with their status history. Duplicate or superseded entities are deactivated, withdrawn from public views, and linked to their replacement.

### Data curation and review process

SysNDD is maintained through continuous manual curation (Figure 1a). New gene-disease associations are added as they appear in the literature. All entities, beginning with those transferred from the previous SysID database, then undergo systematic re-review by experts of the European Reference Network for Rare Malformation Syndromes, Intellectual and Other Neurodevelopmental Disorders (ERN-ITHACA; see Web Resources). During re-review, expert curators reassess evidence categories against current evidence, update HPO phenotype annotations to reflect new clinical data, verify inheritance modes, and add references from recent case reports and large-scale studies. Detailed curation criteria and re-review rules are available in the online SysNDD documentation (see Web Resources). A submitted review becomes public only after approval. Curators in senior roles may approve their own submissions; all other submissions require approval by a second curator.

### The SysNDD toolkit

SysNDD is freely accessible online (Figure 1d; see Web Resources). The web application offers tabular views of entities, genes, phenotypes, and diagnostic panels, with search, filter, and download functions, and a Gene Networks view for exploring the functional clusters and their enrichment tables (Supplemental Methods). Role-gated pages support expert re-review, curator approval, and administration; their access and data-control boundaries are summarized in Table S2. Software can query the same records through the documented public API, and artificial intelligence (AI) assistants can retrieve and cite them through a read-only Model Context Protocol interface. SysNDD evidence categories also serve as training labels for NDDScore, an exploratory genome-wide prediction layer developed separately from curation. NDDScore predicts NDD association, inheritance, and phenotypes for protein-coding genes without altering curated evidence (Table S3).

### Phenotype clustering and functional analyses

To address the fourth question, we clustered the phenotype and functional axes separately and then tested their overlap. Phenotype clustering was performed on Definitive entities with HPO annotations, each encoded by the presence or absence of the 38 HPO phenotype categories. Categories present in fewer than 5% or more than 95% of these entities barely distinguish one entity from another and were excluded (prevalence filtering), leaving 30 HPO categories. The retained binary categories were reduced by multiple correspondence analysis and clustered by hierarchical clustering on principal components^25,26^. The number of clusters was set to the value of *k* at which the step from *k* − 1 to *k* clusters gave the largest proportional drop in within-cluster inertia.

Functional clustering was based on the STRING v11.5 functional-association network^27^, whose separately scored evidence channels can be excluded individually. To limit literature co-mention bias, we excluded the text-mining channel and applied weighted Leiden community detection^28^ to an experimental-plus-database network. Phenotype clusters are defined over gene-disease entities, whereas functional clusters are defined over genes. Correspondence was therefore tested at gene level: each gene was counted once per phenotype cluster, regardless of how many of its entities fell in that cluster, and only genes present in both clusterings were included. For each pair of a phenotype cluster and a functional cluster, we tested whether they share more or fewer genes than expected by chance, using a two-sided Fisher exact test with Benjamini-Hochberg correction across all pairs and a size-preserving permutation test as a robustness check. Overlap is reported as the Jaccard index, the share of genes common to both clusters among all genes in either^29^. Cluster stability, separation, and dependence on curated STRING annotations were evaluated as described in Supplemental Methods. Full parameter settings, confidence-interval procedures, and sensitivity analyses, including a Definitive-only membership check (Table S4), are provided in Supplemental Methods.

### Database comparison and gene count estimation

We compared SysNDD with seven other NDD gene resources: PanelApp, Gene2Phenotype, OMIM (restricted to entries annotated with the HPO term Neurodevelopmental abnormality, HP:0012759, or a descendant; “OMIM NDD”), Radboudumc ID, OrphaID, SFARI Gene, and Geisinger NDD GeneHub^10–13,30,31^. Resource records were mapped to HGNC gene symbols and onto SysNDD’s four evidence categories with a conversion table (the crosswalk), which makes resources with different grading scales, and ungraded gene lists, comparable. Genes that the crosswalk places in the top category, including all genes from ungraded lists, are termed crosswalk-Definitive. The compared records reflect the shared access date of September 21, 2026. The resulting gene-by-resource table and the crosswalk are browsable at https://sysndd.dbmr.unibe.ch/CurationComparisons. Pairwise cosine similarities between binary gene-membership vectors were converted to cosine distances and clustered by average linkage. Completeness was measured as the coverage gap: the share of consensus genes missing from a resource when that resource is excluded from the consensus. As an exploratory analysis, we treated the compared resources as repeated samples from a single gene pool and applied the Chao2 richness estimator^32^ to project the total number of NDD-associated genes with its analytic 95% confidence interval. Sensitivity analyses left out one database at a time and, separately, restricted the input to crosswalk-Definitive genes. Because resource scope and curation networks overlap, the resources are not independent samples, and each projection is descriptive, not a formal lower bound; full methods and sensitivity analyses are provided in Supplemental Methods.

## Results

### A growing catalog in which recessive genes outnumber dominant genes

SysNDD currently catalogs 4,275 gene-inheritance-disease entities (one entity per gene, mode of inheritance, and disease) across 3,273 genes. Of these, 3,758 entities across 3,271 genes are associated with NDDs (accessed September 21, 2026); the other 517 entities record non-NDD disorders, and 2 genes have only such entities. Regarding evidence level for NDD, 2,043 entities are classified as Definitive (1,838 genes; 1,162 autosomal recessive, 698 autosomal dominant, 170 X-linked, and 13 with other inheritance modes), 226 as Moderate (179 genes), and 1,479 as Limited (1,248 genes), with 7 Refuted and 3 uncategorized associations (Figure 1c; inheritance breakdown by evidence category in Figure 2).

**Figure 2:**
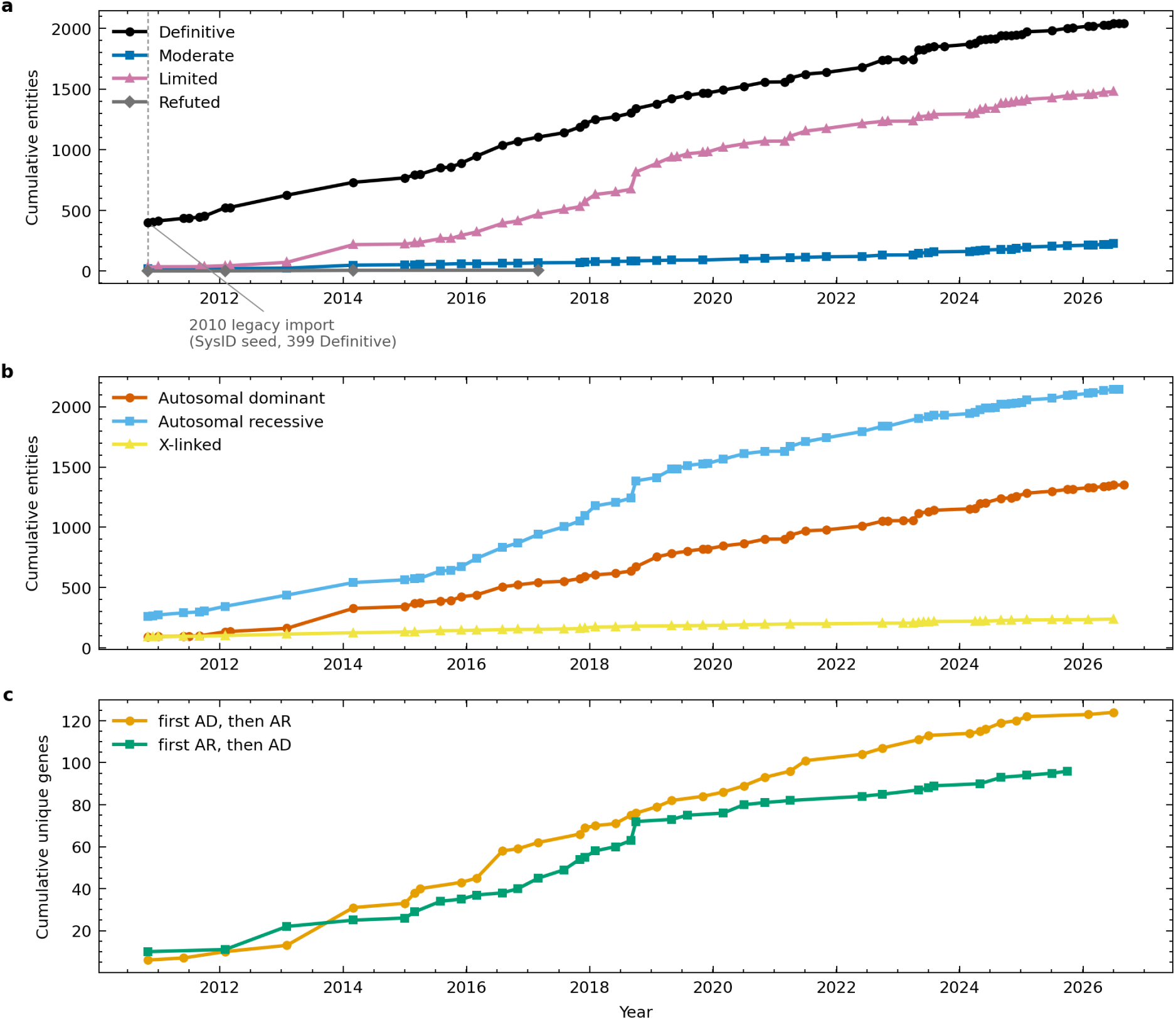
Cumulative gene-disease associations in SysID/SysNDD from 2010 to present. **a**, Cumulative number of gene-disease entities over time, grouped by evidence category (Definitive, Moderate, Limited, or Refuted). Lines represent the running total of entities at each time point at which new entries were added. **b**, Cumulative number of gene-disease entities over time, grouped by inheritance pattern. A gene may be represented by multiple entities with different inheritance patterns. By evidence category, 2,014 genes have a recessive NDD association (1,094 Definitive, 97 Moderate, 822 Limited), 1,262 have a dominant association (639 Definitive, 90 Moderate, 528 Limited), and 200 have an X-linked association (140 Definitive, 12 Moderate, 46 Limited); each gene is counted once per inheritance pattern at the strongest category it reaches for that pattern, and the few genes outside these three categories hold only Refuted or uncategorized assertions for that pattern. **c**, Cumulative number of unique genes associated with both autosomal dominant (AD) and autosomal recessive (AR) NDDs. “First AD, then AR” denotes genes first associated with an AD NDD that later acquired a second, AR association, while “first AR, then AD” denotes genes first associated with an AR NDD that later acquired a second, AD association. Currently, 220 genes have both AD and AR associations, with 124 in the first-AD-then-AR group and 96 in the first-AR-then-AD group. Data: SysNDD API v0.38.1 (timeline accessed September 21, 2026).

Temporal trend analysis (Figure 2; Figure S1) shows substantial growth for both autosomal recessive and autosomal dominant entities. Genes with recessive NDD associations outnumber those with dominant associations (2,014 versus 1,262), and 200 genes have X-linked NDD associations. A total of 220 autosomal genes are associated with both dominant and recessive NDDs (Figure 2c), 45 with Definitive evidence for both entities. Of these dual-inheritance genes, 124 were first associated with a dominant NDD and later gained a recessive association, whereas 96 followed the reverse order. In *SLC2A1* (GLUT1 deficiency syndrome; MIM: 606777) and *DNM1* (developmental and epileptic encephalopathy 31A and 31B; MIM: 616346, 620352), dominant and recessive variants cause the same or allelic disorders. In contrast, in *ACOX1*, for example, they cause distinct diseases: recessive peroxisomal acyl-CoA oxidase deficiency (MIM: 264470) and dominant, gain-of-function-associated Mitchell syndrome (MIM: 618960). X-linked NDDs fit poorly into a dominant/recessive dichotomy anyway, because their manifestation in either or both sexes depends on the gene, the variant mechanism, and X-inactivation^33^. Most X-chromosomal NDD entities in SysNDD therefore lack this attribution: of 238 such entities, 160 are recorded as “X-linked other inheritance”, compared to distinct classification as X-linked recessive inheritance for 61 and X-linked dominant inheritance for 17 entities, respectively. The X chromosome carries relatively few genes, and X-linked NDDs often segregate in families, thus allowing disease gene identification by linkage analysis and Sanger sequencing screens already before high-throughput sequencing^34^. Many X-linked NDD genes were therefore known before exome sequencing, which explains why X-linked entities have grown only modestly in recent years (Figure 2b).

Gene-disease associations in SysNDD, as well as phenotypic and genotypic annotations, are based on 4,932 peer-reviewed publications from 1966 to 2026 (Figure S2). Annual counts of publications used for curation rose steeply through the 2010s, with the broad adoption of exome and genome sequencing in clinical genetics, and peaked at 496 publications in 2021. Counts for the most recent years (215 publications in 2025) are still incomplete, because a publication is counted only once it has been used to curate an entity. Ranked by the number of publications (Figure S2b), the leading sources are the American Journal of Human Genetics (680), GeneReviews (395 chapters, linked as review literature rather than primary references), and the American Journal of Medical Genetics Part A (342).

### SysNDD overlaps with, but is not subsumed by, other NDD gene resources

Of the 3,271 NDD-associated genes in SysNDD, 83.1% are also listed by at least one of seven other major curation efforts, whereas 552 genes with positive SysNDD evidence are listed by none of them (21 Definitive, 23 Moderate, and 508 Limited; Figure 3a). Among broadly scoped databases filtered for neurodevelopmental phenotypes, PanelApp ID has the greatest gene overlap with SysNDD (2,105 genes), followed by OMIM NDD (1,975 genes) and Gene2Phenotype DD (1,907 genes). The smaller, manually curated cross-disorder Geisinger NDD GeneHub shares 1,241 of its 1,576 genes with SysNDD. Gene sets focused on ID or autism overlap with SysNDD to varying degrees: Radboudumc ID shares 1,943 of its 2,010 genes, OrphaID (the Orphanet ID gene list) 1,300 of 1,561, and SFARI 792 of 1,274. The portal states that this gene list was curated in partnership with ERN-ITHACA and SysID^30^, SysNDD’s predecessor, so the SysNDD–OrphaID overlap partly reflects shared curation provenance rather than independent agreement.

**Figure 3:**
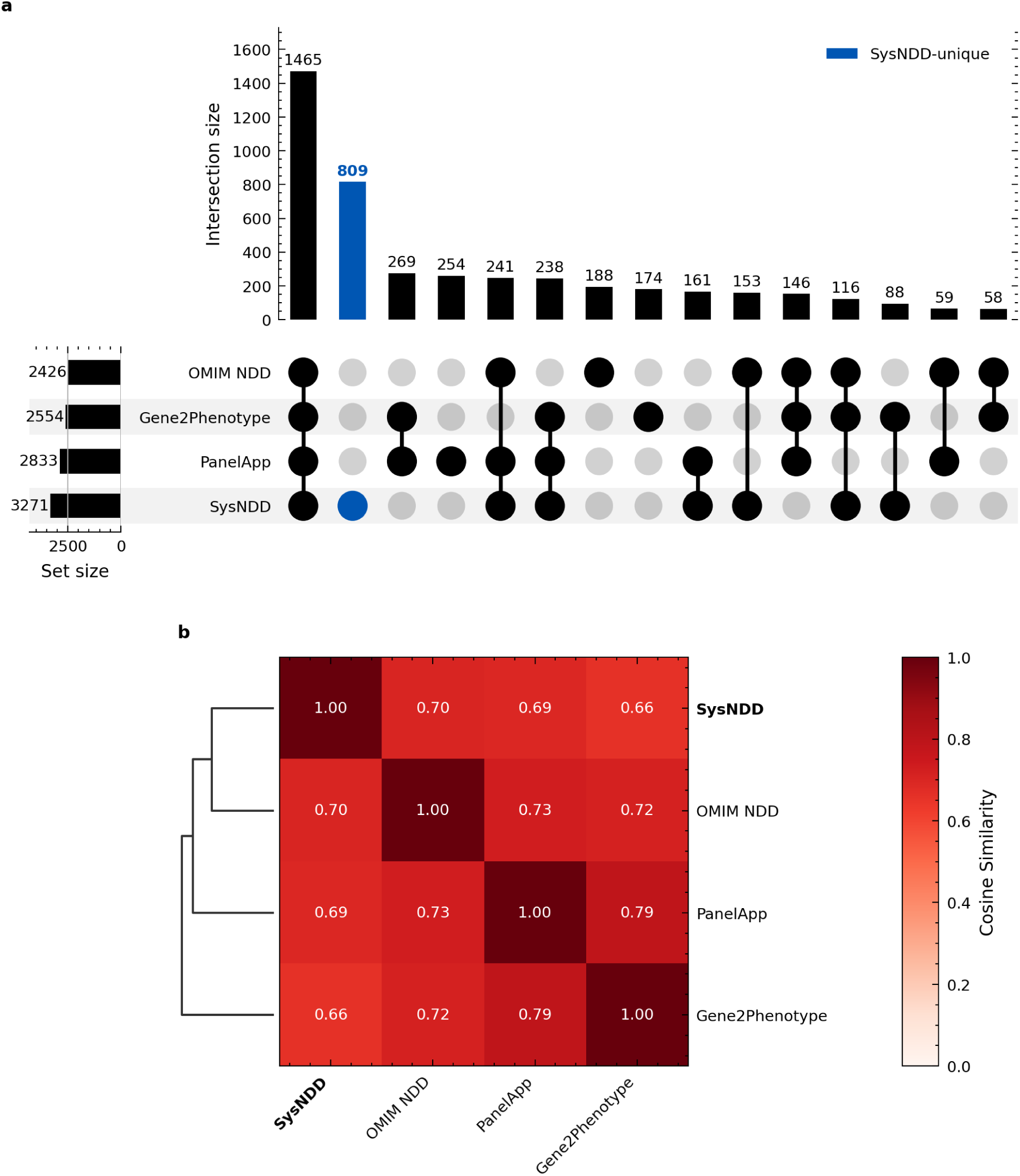
Comparison of SysNDD with other neurodevelopmental disorder gene databases. **(a)** UpSet plot^35^ showing gene overlaps between the four largest NDD databases: SysNDD (3,271 genes), PanelApp (2,833), Gene2Phenotype (2,554), and OMIM NDD (2,426). Bars are ordered by intersection size. Among the four resources shown, 809 genes are unique to SysNDD; across all eight analyzed resources, 552 positive-evidence SysNDD genes are absent from all seven comparators (21 Definitive, 23 Moderate, and 508 Limited). All four resources share 1,465 genes. **(b)** Cosine similarity heatmap with hierarchical clustering showing relationships between the four databases. Databases with more similar gene content cluster together. Among these four databases, SysNDD is most similar to OMIM NDD, while the closest pair overall is PanelApp and Gene2Phenotype. Data: SysNDD API v0.38.1 (accessed September 21, 2026).

The most similar gene lists are those of Gene2Phenotype DD and PanelApp ID (cosine similarity 0.79, where 1 means identical lists; Figure 3b; all eight resources in Figure S3b), consistent with shared curation networks. SFARI is the least similar (similarity ∼ 0.3–0.43), reflecting its focus on autism, a specific NDD subset, whereas the cross-disorder Geisinger NDD GeneHub is more similar (∼ 0.43–0.58), consistent with its broad neurodevelopmental scope. Across all eight compared resources, SysNDD is most similar to the Radboudumc ID gene list, compatible with, but not proof of, shared curation traditions and collaborative networks.

The Chao2 richness estimator, which infers how many genes remain unlisted from how many appear in only one or two resources, projected about 6,865 NDD genes in total (95% CI 6,639–7,126), against 5,166 observed across these resources (Figures S4 and S5). Of the genes listed by at least four other resources, only 4.8% were absent from SysNDD (its coverage gap), a smaller gap than for any other compared resource (Figure S6). When only crosswalk-Definitive genes are counted (genes in the top category after mapping every resource onto the SysNDD scale), the ranking reverses: Radboudumc ID (2.2%) and OMIM NDD (5.6%) have smaller four-database coverage gaps than SysNDD (6.8%). Most genes behind this gap (80 of 92) are listed by SysNDD at a lower category, among them *SPAST* and *NPC1*, whose degenerative disorders involve variable or acquired cognitive impairment; only 12 are not listed as NDD genes. The Definitive-only gap therefore mainly reflects stricter grading rather than missing genes.

Beyond coverage, we assessed how well SysNDD’s evidence categories agree with those of external gene-disease validity resources for co-curated gene-disease pairs, collapsed to SysNDD’s four-category scale (Table S1; Figure S7). Of the 565 gene-disease pairs curated by both SysNDD and ClinGen, the prespecified primary comparator, the two resources assigned the same evidence category to 72% and the same or an adjacent category to 84.6%. Weighted tier agreement was 85.5% (Gwet’s AC2 0.806, 95% CI 0.774–0.839). Cohen’s *κ* was nevertheless low (0.191), because most co-curated pairs were rated Definitive by both resources, and *κ* penalizes this skew even when agreement is high (Table S5). Gene2Phenotype and Genomics England PanelApp showed similar agreement (AC2 0.833 and 0.837).

We then categorized all 158 SysNDD–ClinGen discordances by evidence type using a large language model that read the primary curation records of both resources (Supplemental Methods; Data S1). Of these, 51.3% were genuine disagreements about the evidence category, and in 77.8% of these SysNDD held the higher evidence category. The rest arose from the comparison rather than from the evidence, chiefly from differing curation dates and disease scope. Of the 44 discordances attributed to curation-date lag, SysNDD retained a lower category after ClinGen had promoted the association in 39; 2 showed the reverse pattern, and the remainder could not be assigned a direction. These lags typically spanned several years (Table S5). Blinded expert re-review of a stratified 60-discordance sample agreed moderately with the model (Cohen’s *κ* = 0.433), so these proportions are model-assisted estimates.

### Recorded phenotypes portray NDDs as multisystem disorders, and variant annotations follow molecular logic

We examined how the 38 annotated HPO phenotype categories are distributed across the 3,758 NDD entities in SysNDD (14,545 total annotations; Figure 4a, Figure S8a). The most frequent categories are “abnormality of the nervous system” (n=1,277), “abnormality of brain morphology” (n=1,173), and “seizures” (n=1,143). “Behavioral abnormality” (n=697) and “abnormality of the eye” (n=674) are also prevalent, while “genitourinary abnormalities” (n=263) and “Progressive” (n=303) illustrate the multisystem and sometimes (neuro)degenerative nature of NDDs. Intellectual disability is recorded not by the base HPO term (HP:0001249) but by five ID severity categories, which reflect the degree of cognitive impairment rather than accompanying phenotypes; “intellectual disability, moderate” is the most frequent (n=1,051; Figure 4a).

**Figure 4:**
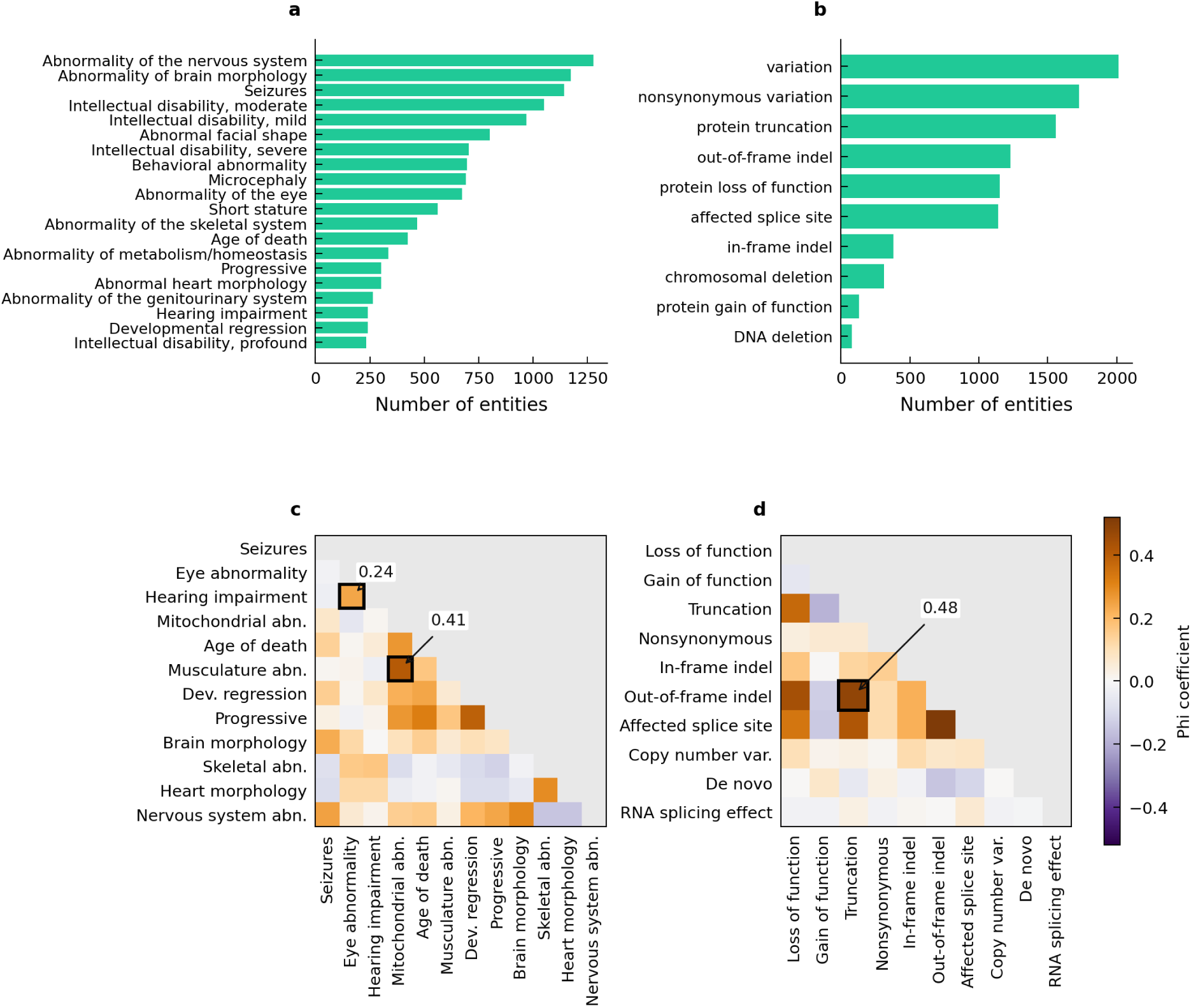
Phenotype and variant type distributions and correlations in SysNDD. **a**, Horizontal bar chart showing the frequency of Human Phenotype Ontology (HPO) phenotype categories across SysNDD entities (top 20 of 38 categories shown; complete distribution in Figure S8). Categories are sorted by entity count in descending order. Abnormality of the nervous system (HP:0000707) is the most frequently annotated phenotype (n=1,277), followed by abnormality of brain morphology (HP:0002011, n=1,173) and seizures (HP:0001250, n=1,143). **b**, Distribution of variant types across entities (top 10 of 40 categories shown), classified using the Variation Ontology (VariO). Among the specific classes, nonsynonymous variation (VariO:0017, n=1,728) and protein truncation (VariO:0015, n=1,558) are the most frequent. **c**, Phenotype correlation heatmap showing pairwise Pearson *r* between binary phenotype-category presence vectors (the phi coefficient) among 12 selected HPO categories. The lower triangle is shown; the unit diagonal and duplicate upper triangle are masked. Highlighted cells indicate the mitochondrial-musculature correlation (*r* =0.41) and the eye-hearing sensory correlation (*r* =0.24). Purple indicates negative and orange positive association; panels **c** and **d** share a symmetric diverging color scale spanning the wider of the two panels’ correlation ranges. Full 38-phenotype matrix in Figure S10. **d**, Variant-consequence correlation heatmap over a curated subset of 10 mechanistically interpretable Variation Ontology (VariO) classes with at least 10 annotated entities. A strong, mechanistically interpretable association (highlighted) links protein truncation / out-of-frame indel (*r* =0.48), consistent with out-of-frame (frameshifting) indels producing truncated proteins. Color encoding and triangular display are as in **c**. Full 40-category variant matrix in Figure S11. HPO and VariO annotations are hierarchical and non-exclusive; the correlations are descriptive, not causal. Data: SysNDD API v0.38.1 (accessed September 21, 2026).

We call an entity syndromic when at least one organ system outside the nervous system is recorded (16 systems, each counted once; Supplemental Methods). Of the 2,017 Definitive NDD entities with at least one recorded phenotype, 1,688 (83.7%) are syndromic by this definition, and 1,310 (64.9%) have two or more organ systems involved. Craniofacial features, abnormal head size, and eye involvement are most frequently annotated. Because micro- and macrocephaly may reflect brain involvement rather than a separate organ system, we repeated the count without head size; the syndromic share then falls to 78.8% (Figure S9). Features not reported in the literature count as absent, so some entities classed as non-syndromic may in fact be syndromic.

SysNDD captures variant consequences and types using 40 VariO categories^22^ (10,072 total annotations; Figure 4b, Figure S8b). Among specific consequences, “nonsynonymous variation” is most prevalent (n=1,728), followed by “protein truncation” (n=1,558), “out-of-frame indel” (n=1,229), explicit “protein loss of function” (n=1,151), and “affected splice site” (n=1,141). “Chromosomal deletion” (n=312), a VariO class distinct from the indel classes, and gain-of-function effects (n=133) are also documented. Because VariO annotations are hierarchical and non-exclusive, these counts describe annotation patterns rather than frequencies of molecular mechanisms. The annotations are curated from the primary literature and maintained separately from the ClinVar^24^ records that SysNDD displays for the same gene, so the two sources may differ.

Phenotype co-occurrence, measured as phi coefficients between the presence of two categories across Definitive entities and reported descriptively (Supplemental Methods), reveals biologically coherent correlations. Figure 4c shows 12 categories selected manually to span neurological, sensory, organ-system, and clinical-course features; the full 38-phenotype matrix is shown in Figure S10. “Abnormality of the mitochondrion” correlates with “musculature abnormality” (r = 0.41) and with “age of death” (r = 0.27), consistent with the severe clinical course of some mitochondrial disorders. Weaker but expected pairings link “abnormality of the eye” with “hearing impairment” (r = 0.24) and “seizures” with “abnormality of brain morphology” (r = 0.23).

Variant-consequence annotations show comparable internal coherence (Figure 4d; full 40-category matrix in Figure S11). Within the curated subset shown in Figure 4d, the strongest positive associations follow the expected molecular logic, from a frameshifting or splice-disrupting change to a truncated, non-functional protein: affected splice site / out-of-frame indel (r = 0.52), protein truncation / out-of-frame indel (r = 0.48), and protein loss of function / protein truncation (r = 0.37). The strongest negative association, though modest, is protein gain of function / protein truncation (r = -0.19), consistent with gain and loss of function being recorded as alternative rather than co-occurring mechanisms. Together, the phenotype annotations portray NDDs as multisystem disorders, and both phenotype and variant annotations show internally coherent patterns, although variant mechanism is captured only coarsely for much of the catalog.

### Genes form distinct molecular clusters, whereas disorders form broad phenotype groups

The two clustering approaches address different questions: phenotype clustering groups gene-disease entities whose clinical presentations resemble one another, whereas functional clustering groups genes whose protein products act together in the cell, irrespective of the associated disorders. Comparing the two partitions tests a long-standing proposal that phenotypically similar disorders often arise from variants in functionally related genes^36^. Genes fell into 18 functional clusters (Figure 5a) and Definitive entities into 3 broad phenotype clusters (PC1–3; Figure 5b; Tables S6 and S7). Functional clusters keep their raw Leiden community numbers, so their labels are not consecutive (e.g., FC69, FC301).

**Figure 5:**
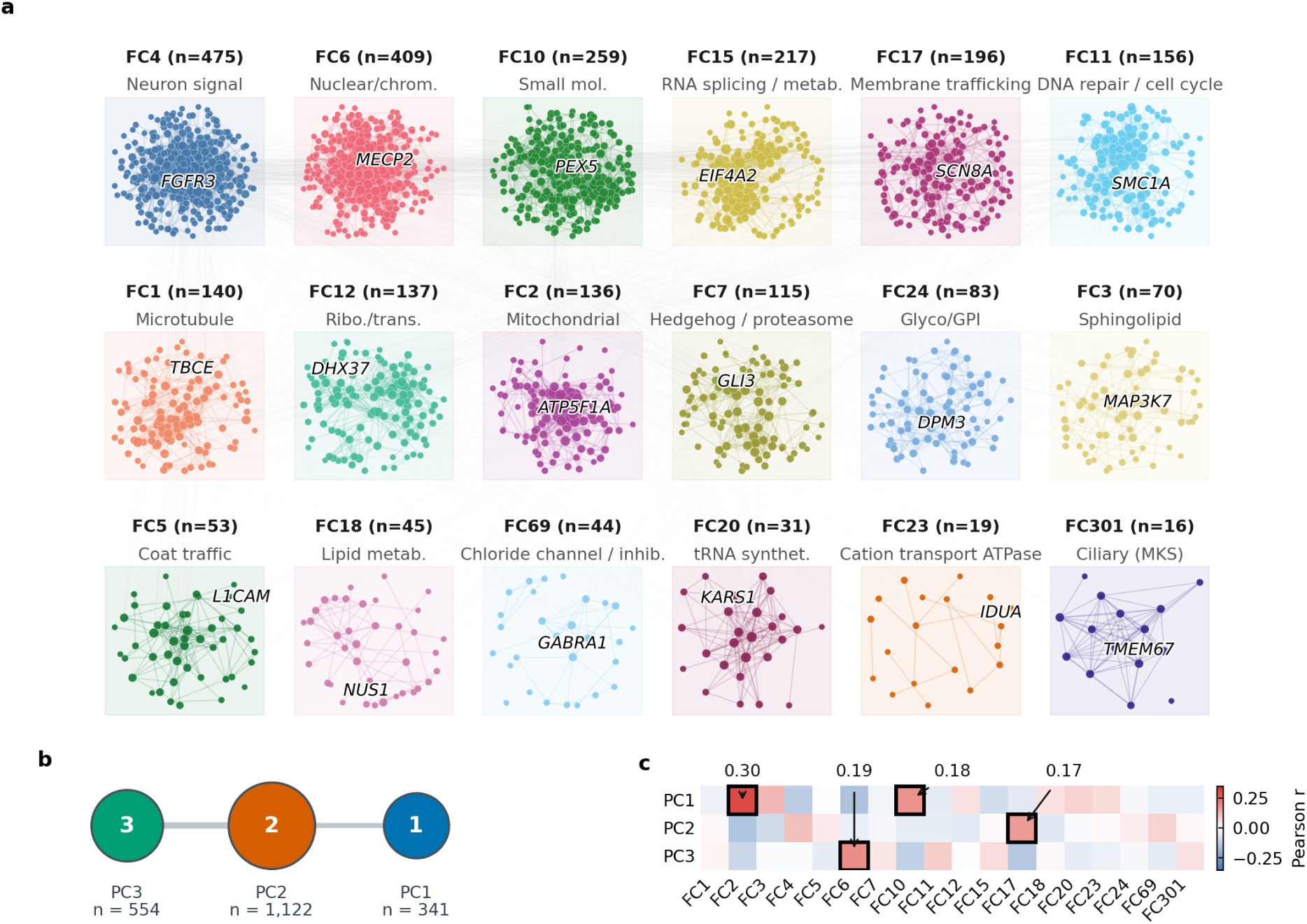
Functional clusters show selective, generally weak correspondence with phenotype clusters. **(a)** STRING v11.5 functional-association network^27^ of NDD-associated genes, partitioned by weighted Leiden community detection into 18 functional clusters. Each deterministic force-directed subnetwork^37^ shows its functional theme, full membership count, and labeled representative gene; nodes are scaled by displayed interaction degree. For legibility, the 2,601 clustered genes are represented by the 10,000 strongest interactions, as a display subset of the graph used for clustering. **(b)** Phenotype clusters from multiple correspondence analysis followed by hierarchical clustering on the prevalence-filtered, category-level HPO representation of Definitive entities: PC1, progressive/metabolic (n=341); PC2, intellectual disability/seizures (n=1,122); and PC3, multisystem malformation (n=554). Node area encodes cluster size, link width encodes the absolute correlation between the two clusters’ memberships, and the layout is not a spatial embedding. **(c)** The PC–FC correlation block is reconstructed directly from the memberships displayed in panels **a** and **b**, using one binary membership bit per gene across the 1,517 genes of the shared gene set. Rows show PC1–PC3 and columns show FC1–FC301 (raw Leiden community numbers, hence non-consecutive); color encodes signed Pearson *r*. Black outlines with coefficient labels mark the four associations discussed in the Results (PC1–FC2, PC1–FC10, PC3–FC6, and PC2–FC17). They are among the 17 of 54 cells significant at q < 0.05; the other significant cells are not outlined. Associations across the block are generally small (median |*r* |=0.05, maximum 0.30). Data: SysNDD analysis release (API v0.38.1; accessed September 21, 2026).

The two axes behave very differently. Genes cluster much more strongly than expected by chance: the modularity of the functional network (0.679) far exceeds that of randomized networks in which every gene keeps its number of interactions (mean 0.22; empirical *p* ≤ 0.005). The phenotype partition is reproducible under resampling (bootstrap-Jaccard 0.935–0.947, where 1 means the same clusters on every resample). It is, however, only weakly separated (mean silhouette 0.183, on a scale where 0 means no separation and 1 perfect separation), so there is no evidence of discrete phenotype modules in this representation. Molecular interaction is thus the sharper organizing axis, although the network partly re-encodes curated pathway annotation that the phenotype side also draws on. Stability of the functional partition varies by cluster: per-cluster bootstrap-Jaccard ranges from 0.326 in FC23 to 0.813 in FC10 (Table S7). Re-clustering on experimental evidence alone, without STRING’s curated-database channel, still gave strong modularity (0.74, z = 206 against the same null), but the resulting clusters agreed only moderately with the combined ones (adjusted Rand index 0.39, where 1 means identical clusters; Figure S12).

The phenotype clusters capture broad clinical axes rather than fine syndromic groups. PC1 (341 entities) gathers progressive, metabolic, and mitochondrial presentations and is enriched for autosomal recessive inheritance; PC2 (1,122 entities) is dominated by intellectual disability, behavioral abnormalities, and seizures; and PC3 (554 entities) is enriched for genitourinary, skeletal, cardiac, facial, and renal abnormalities and short stature. PC2 and PC3 are both enriched for autosomal dominant inheritance. All of them are weakly separated (per-cluster silhouettes 0.05–0.251) in this HPO representation. The partition also depends on how missing annotations are treated: it is not reproduced when unrecorded phenotypes are treated as unknown rather than absent (adjusted Rand index near zero). The partition was largely retained when the intellectual disability severity grades were removed or prevalence filtering was omitted (adjusted Rand index 0.75 and 0.83; Supplemental Methods). The 18 text-mining-free functional clusters, by contrast, resolve recognizable molecular themes.

Representative genes include *NDUFS1* and *ACAD9* (mitochondrial function); *ARID1B*, *CHD7*, and *KMT2D* (chromatin regulation); *GRIN1* and *SYNGAP1* (synaptic signaling); *CEP290* (ciliary transition zone); and *PIGA* and *PIGN* (glycosylation). All themes are listed in Table S7. A further 550 NDD genes have no interaction above the score threshold once the text-mining channel is excluded and therefore belong to no cluster. The molecular composition of the phenotype clusters, which their construction does not use, is tested next.

### Clinical groups correspond to molecular clusters only selectively

We tested whether phenotype clusters align with functional clusters across 54 pairwise comparisons (Figure 5c; Tables S6 and S7) among the 1,517 genes present in both clusterings (1,823 genes carry a phenotype cluster and 2,601 a functional cluster). Of the 54 cells, 17 were significant (q < 0.05; 9 enrichments and 8 depletions), but the overlaps were modest. The most readily interpretable coupling was the near-definitional PC1–FC2 link between the progressive/metabolic phenotype cluster and the mitochondrial cluster (OR=11.05, 95% CI 6.83 to 17.87, Jaccard=0.182, q=<0.001; full test statistics in Data S1). An odds ratio of this size means that progressive/metabolic-cluster genes are many times over-represented in the mitochondrial cluster, whereas the low Jaccard index means that the two groupings share only a minority of their genes.

Beyond this definitional axis, the progressive/metabolic cluster was also enriched for the small-molecule-metabolism cluster (PC1–FC10; OR=3.37, Jaccard=0.165). The multisystem-malformation cluster was enriched for the nuclear/chromatin cluster (PC3–FC6; OR=2.77, Jaccard=0.216), consistent with chromatin regulators such as *ARID1B*, *CHD7*, and *KMT2D* in that cluster. Likewise, the intellectual disability/seizure cluster was enriched for the neuronal-signaling and membrane-trafficking clusters (PC2–FC4, OR=1.82; PC2–FC17, OR=4.91). Phenotype clusters therefore do not map one-to-one onto molecular modules. Of the couplings highlighted in Figure 5c, 3 persist after removing STRING’s curated-database channel (Supplemental Methods), whereas the small-molecule-metabolism link remains specific to the combined network.

Because this comparison inherits the weak phenotype partition, we also tested each functional cluster against the 30 active phenotype categories (Figure S13). Of the 540 cells, 110 deviated from expectation (q < 0.05), including kidney abnormalities in the ciliary cluster and microcephaly in the DNA-repair cluster. Overall, the two axes correspond only partially: significant enrichments or depletions occur in a minority of cluster pairs, individual overlaps are modest, and the most prominent coupling, PC1–FC2, is near-definitional.

## Discussion

SysNDD represents each neurodevelopmental gene-disease association as a versioned gene-inheritance-disease entity with a recorded evidence category and supporting publications. This structure lets clinicians and researchers query individual associations, assemble or audit diagnostic panels, analyze the catalog systematically, and trace when and why an assessment changed.

Curation in SysNDD began with the transition from SysID on May 7, 2022. Since then, the database has added 820 gene-disease associations (762 unique genes), averaging 187 associations and 174 genes per year. Over the same period, 689 re-reviews (systematic expert reassessments, beginning with entries transferred from SysID) were completed, and the annotations of 1,496 entities and the status of 837 entities were updated, including for dual-inheritance patterns and phenotypic expansions. These activities show that continuous curation involves both expanding the catalog and reassessing existing associations.

The clinically most relevant differences between resources involve Definitive and Moderate evidence level categorization. SysNDD includes 21 Definitive and 23 Moderate genes absent from every other comparison resource. These higher-confidence associations illustrate the complementary coverage of expert-curated resources^8^. However, their absence from another resource does not necessarily indicate disagreement, as resources differ in scope, inclusion criteria, evidence frameworks, and update schedules. The 508 SysNDD-only Limited genes include early and historical candidate associations that require continuing re-review; where current criteria are no longer met, they are demoted or reclassified as Refuted rather than removed. A continuously maintained resource can promote, demote, or refute associations as evidence changes^17^, whereas a published list begins aging at publication. Continuous maintenance does not remove reassessment lag, however: in most date-lag discordances with ClinGen, SysNDD was behind, which argues for periodic cross-resource review.

The distribution of inheritance modes has implications for diagnostic strategies and genetic counseling. Although autosomal dominant NDDs associated with *de novo* variants predominate in non-consanguineous diagnostic cohorts^1^, autosomal recessive entities outnumber autosomal dominant entities in SysNDD, consistent with the even more pronounced genetic heterogeneity of recessive NDDs^38^. After proband-only sequencing, recessive diagnoses may require parental testing or long-read data to phase variants and establish compound heterozygosity. The substantial fraction of *de novo* dominant diagnoses, in turn, helps explain the higher diagnostic yield of trio-based than proband-only sequencing^1,39^. In the 220 genes with both dominant and recessive entities, a heterozygous variant may be causal or indicate only carrier status, so interpretation and counseling must be mode-specific. X-linked labels are similarly imprecise. Females with heterozygous pathogenic variants in genes associated with only X-linked recessive NDDs may be symptomatic, and female predominance in some X-linked dominant disorders may reflect male lethality^33,40^.

Inheritance mode and variant mechanism must also be interpreted together. Loss-of-function variants contribute to both recessive and dominant NDDs, through biallelic loss of function in recessive disorders and haploinsufficiency in many dominant disorders. Common examples of disease genes with haploinsufficiency mechanisms include *ARID1B* (Coffin-Siris syndrome 1; MIM: 135900), *ANKRD11* (KBG syndrome; MIM: 148050), and *SYNGAP1* (intellectual developmental disorder, autosomal dominant 5; MIM: 612621), whereas other dominant NDD genes act through gain-of-function, dominant-negative, or other mechanisms^41,42^, and in some genes, such as *ACOX1*, different mechanisms cause distinct disorders. Copy-number changes and other structural variants add another layer and may be detected by exome-based dosage analysis^43^, genome sequencing, or targeted assays. SysNDD records VariO classes for each entity, allowing molecular mechanism to be considered alongside inheritance and clinical interpretation.

In the current catalog, molecular modules are well separated, whereas disorder-level HPO profiles form broad, overlapping groups. Part of this asymmetry reflects resolution: each disorder is described by 30 binary phenotype categories, whereas each gene is placed by weighted interactions with its partners. Aggregation of phenotypes across individuals within a disorder and incomplete phenotype annotation may further limit correspondence between phenotypic and molecular groupings, as anticipated by Kochinke et al. for sparse disorder-level phenotyping^18^. These clusters and their correspondence are descriptive. They neither define a phenotype taxonomy nor establish causal relationships between molecular modules and phenotypes. For now, molecular modules offer the more robust grouping of NDD genes, and phenotype groups are best read as broad clinical context until patient-level phenotypes allow a finer comparison.

SysNDD’s distinct contribution lies in its independent primary curation of versioned gene-inheritance-disease entities, including separate entities for distinct disease associations or modes of inheritance of the same gene. This complements GenCC’s harmonization of assertions from member groups^16^. The structured annotations associated with these entities support NDD-scale phenotype-function analyses: shared pathways can underlie overlapping, pleiotropic phenotypes^44,45^, and NDD-associated genes converge in co-expression and interaction modules^46,47^.

SysNDD evidence categories also provide the training labels for NDDScore, the exploratory genome-wide prediction layer: genes with Definitive or Moderate associations form its positive reference set. On held-out test data, its association model achieved an area under the receiver-operating characteristic curve of 0.89 (0.5 is chance and 1 perfect ranking). Applied to the 16,088 genes not curated in SysNDD, it ranked *MAU2*, the obligate partner of *NIPBL* in cohesin loading, highest. *MAU2* was not part of the training data and was only later included in SysNDD, after a recent report established it as a new Cornelia de Lange syndrome gene^48^, consistent with the predicted autosomal dominant inheritance and phenotypes. This is a post hoc observation, not an independent validation.

Disorder-level HPO annotations do not retain patient-level detail and depend on what the literature reports and curators capture, so they cannot show how features are distributed across individuals. Phenotypes recorded per patient in a standardized, machine-readable form could test whether the same groupings emerge in individuals^49^. The functional analyses have a related limit: they include only genes with interaction evidence beyond co-mention in the literature, and because the network partly draws on known pathway annotations, its modules and their phenotype associations are not fully independent of existing knowledge.

Because entities are versioned and record phenotype, inheritance, and variant consequence in one structure, SysNDD supports analyses beyond a pure gene list. Genes retrieved by a combination of phenotype categories can be intersected with functional modules to nominate mechanisms for experimental follow-up, as proposed for intellectual disability modules^18^; the recovery of the known kidney involvement of ciliopathy genes (Figure S13) shows that such intersections return expected biology. These queries generate hypotheses and inherit the limits of literature-derived, disorder-level annotation. Finally, both Chao2 projections indicate that a substantial share of NDD genes remains unlisted; continued curation will test them prospectively, as it will the NDDScore rankings.

## Supporting information

Data S1 analysis tables

## Data and code availability

The SysNDD database is freely accessible at https://sysndd.dbmr.unibe.ch (permanent redirect: https://sysndd.org). All data are available through the public API at https://sysndd.dbmr.unibe.ch/API with full OpenAPI/Swagger documentation. The SysNDD web application source is publicly available at https://github.com/berntpopp/sysndd, and the key analysis software and packages are listed in Table S8. Database content is released under the Creative Commons Attribution 4.0 International (CC BY 4.0) license (https://creativecommons.org/licenses/by/4.0/).

This paper analyzes existing, publicly available data: the SysNDD records and analysis releases served by API v0.38.1 on the access dates given in the figure legends and Supplemental Methods. Source URLs and access dates are listed in Table S9. The underlying analysis tables are provided as Data S1, a workbook with one worksheet per table and a README sheet describing each. The separately developed NDDScore prediction release is freely available at https://sysndd.dbmr.unibe.ch/NDDScore and archived at Zenodo (concept DOI 10.5281/zenodo.20258026).

## Supplemental information

Supplemental information (Supplemental Methods, Figures S1–S13, Tables S1–S9, Note S1, and Data S1) accompanies this article.

## Acknowledgments

C.Z. and B.P. were supported by grants from the Deutsche Forschungsgemeinschaft (DFG; ZW 184-6/1, PO 2366/2-1). A.S. was supported by the Netherlands Organisation for Scientific Research (ZonMw Vici grant 09150181910022 and a ZonMw Vidi grant). This study has been supported by European Reference Network on Rare Congenital Malformations and Rare Intellectual Disability (ERN-ITHACA). ERN-ITHACA is funded by the European Union, under the grant agreement No. 101156387.

## Author contributions

B.P.: Conceptualization, Data curation, Formal Analysis, Funding acquisition, Investigation, Methodology, Project administration, Software, Visualization, Writing - original draft, Writing - review & editing. S.F.: Data curation, Investigation, Validation, Writing - review & editing. M.F.A.: Data curation, Investigation, Validation, Writing - review & editing. H.V.E.: Data curation, Investigation, Writing - review & editing. A.C.: Data curation, Investigation, Writing - review & editing. N.C.B.: Data curation, Investigation, Writing - review & editing. F.H.: Data curation, Investigation, Writing - review & editing. S.G.: Data curation, Investigation, Writing - review & editing. T.K.: Data curation, Investigation, Writing - review & editing. A.S.: Conceptualization, Writing - review & editing. Z.T.: Data curation, Investigation, Writing - review & editing. A.V.: Funding acquisition, Writing - review & editing. C.Z.: Conceptualization, Data curation, Formal Analysis, Funding acquisition, Investigation, Methodology, Project administration, Supervision, Writing - original draft, Writing - review & editing.

## Web resources

SysNDD database, https://sysndd.dbmr.unibe.ch

SysNDD redirect URL, https://sysndd.org

SysNDD API documentation, https://sysndd.dbmr.unibe.ch/API

SysNDD Model Context Protocol interface, https://sysndd.dbmr.unibe.ch/mcp

SysNDD curation criteria, https://berntpopp.github.io/sysndd/05-curation-criteria.html

ERN-ITHACA, https://ern-ithaca.eu

## Declaration of generative AI and AI-assisted technologies in the manuscript preparation process

During the preparation of this work the authors used Anthropic Claude (Opus 4.8, Opus 5, Fable 5, and Fable 5.1) and OpenAI GPT-5.5 and GPT-5.6 to support language refinement, clarity of expression, and consistency of formatting in the manuscript. After using these tools, the authors reviewed and edited the content as needed and take full responsibility for the content of the publication. Use of large language models as part of the research process is described in the Results and Supplemental Methods for the automated adjudication of curation discordances and in the Supplemental Methods for the generation of functional-cluster pathway summaries and phenotype-cluster summaries.

