## Supplemental Information for "SysNDD: A Systematic Database for Neurodevelopmental Disorders"

for

### Supplemental Figures

#### Figure S1 | Gene-Disease Additions per Year across the SysID and SysNDD Eras

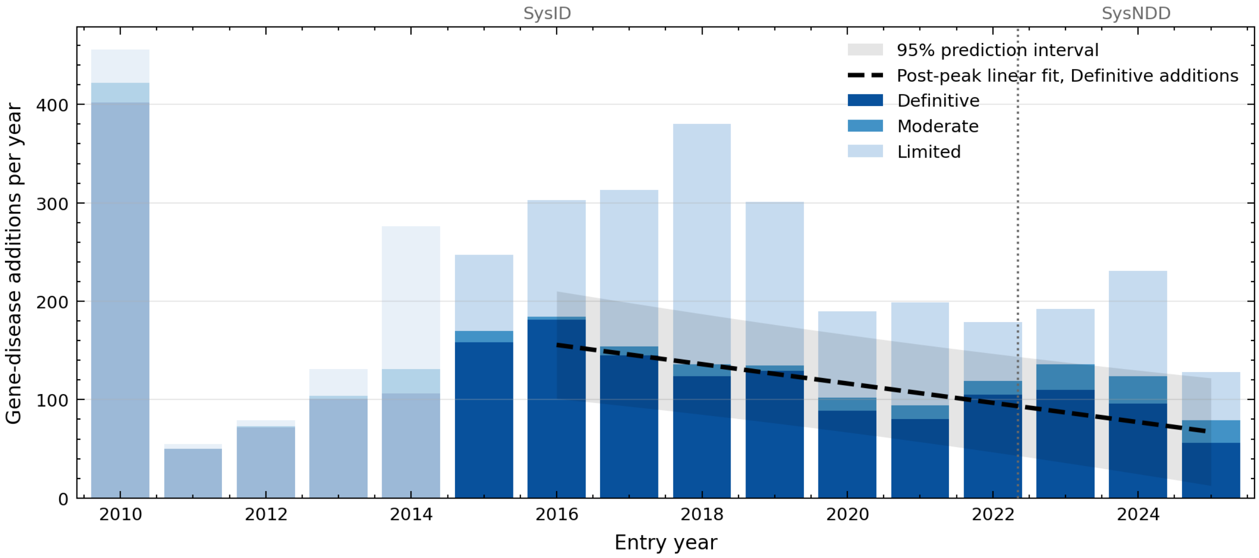

**Figure S1. Gene-disease additions per entry year across the SysID and SysNDD curation eras.** Stacked bars show gene-disease entities added per entry year, shaded by evidence category (dark = Definitive, mid = Moderate, light = Limited); Refuted entities are omitted. Entities curated in the predecessor SysID database were transferred to SysNDD with their original entry dates, so the series spans both systems. SysID was seeded in November 2010 with an initial set of 399 Definitive entities and updated in occasional batches before 2015 (faded bars) and several times each year from then on. Curation in SysNDD itself began on May 7, 2022 (vertical dotted line; eras labeled above the panel). The dashed line is a linear fit to Definitive additions from their 2016 peak (the highest year from 2015 onward) through 2025 (n = 10 years, $R^2$ = 0.70, p = 0.002, slope -9.8 entities per year). The faded wedge is its 95% prediction interval. The fit spans both eras and is descriptive, not a forecast of field-wide gene discovery. The current, incomplete year is omitted. Data: SysNDD API v0.38.1 (timeline accessed September 21, 2026).

### Figure S2 | Publication Statistics

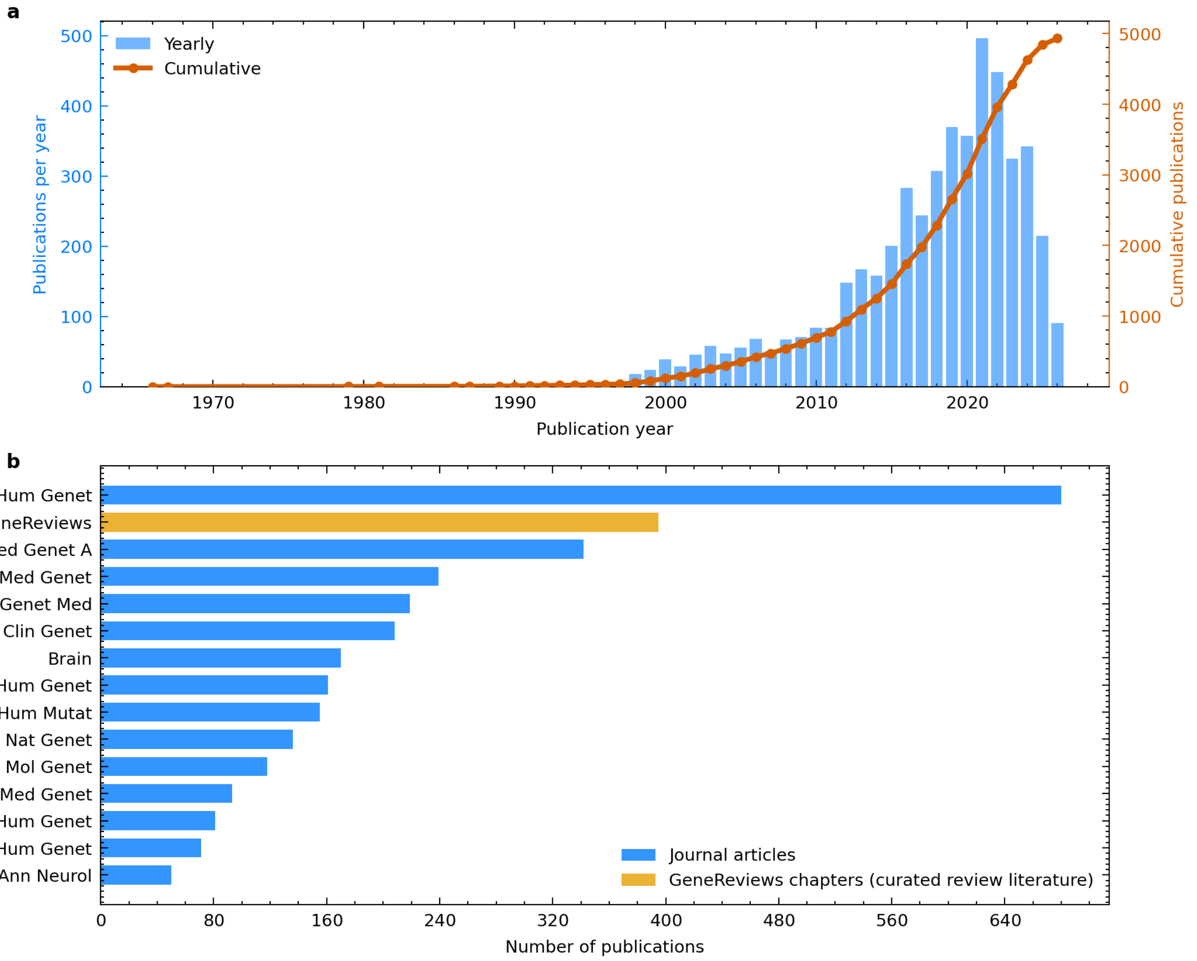

**Figure S2. Publication statistics underlying SysNDD gene-disease curation.** (**a**) Publications used for curation by year of publication, from 1966 to 2026. Publications are placed at their publication year, not the curation date, so the series reaches back decades before SysID[1] or SysNDD existed; Figure 2, by contrast, dates entities by database entry. Bars show publications per year; the line shows the cumulative total. Annual counts rose through the 2010s, peaked at 496 in 2021, and stood at 215 in 2025; the most recent years are incomplete because each publication is counted only once it has been used for curation (4,932 publications in total; five-year average of 365 per year). (**b**) Top 15 sources by publication count. GeneReviews (orange) is not a journal but the NCBI Bookshelf collection of expert-authored disease chapters; its 395 chapters are linked through a dedicated review-literature field, not as primary references. Among journals (blue), the American Journal of Human Genetics is the leading source with 680 publications, followed by the American Journal of Medical Genetics Part A (342) and the Journal of Medical Genetics (239). The publications are drawn from 412 distinct sources. Data: SysNDD API v0.38.1 (accessed September 21, 2026).

### Figure S3 | Extended Database Comparison

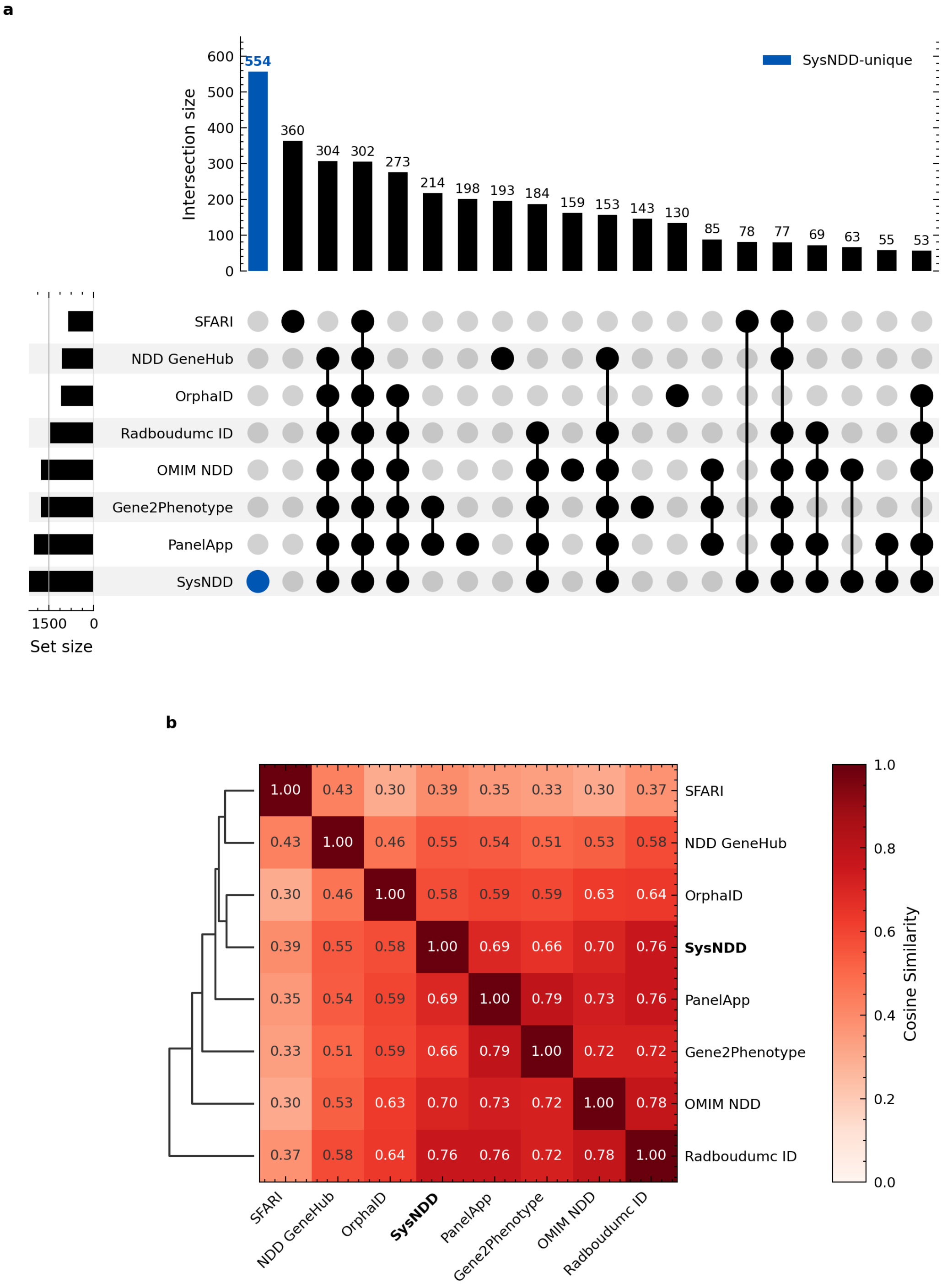

**Figure S3. Extended comparison of SysNDD with seven other neurodevelopmental disorder gene databases.** (**a**) UpSet plot[2] of gene overlaps between all eight compared databases: SysNDD, PanelApp, Gene2Phenotype, OMIM NDD, Radboudumc ID, OrphaID, SFARI, and Geisinger NDD GeneHub. Bars are ordered by intersection size; smaller intersections are omitted for legibility. A gene belongs to the set for a resource when that resource lists it in any category, including Refuted. The SysNDD-only bar (554 genes) therefore includes 3 genes that SysNDD itself lists as Refuted and excludes 1 that another resource lists only as Refuted, which accounts for the 552 positive-evidence SysNDD-only genes reported with Figure 3. (**b**) Cosine similarity heatmap with hierarchical clustering of all eight databases; databases with more similar gene content cluster together. Data:

SysNDD API v0.38.1 (accessed September 21, 2026).

#### Figure S4 | Rarefaction across NDD Gene Resources

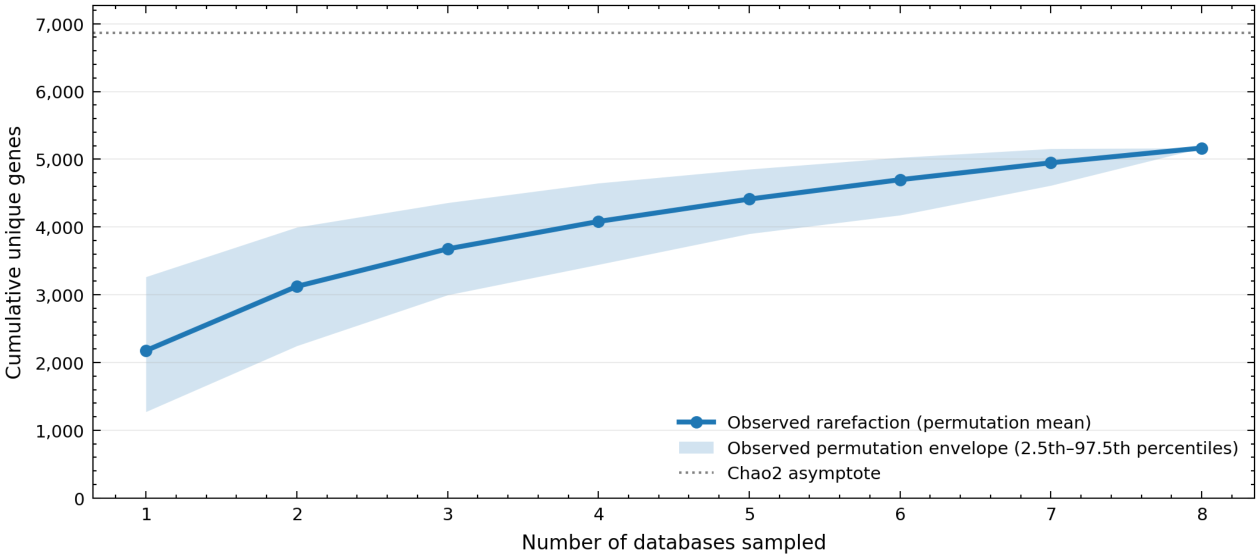

**Figure S4. Rarefaction across NDD gene resources.** Rarefaction shows how the number of distinct genes grows as databases are added one at a time; a flattening curve indicates that further databases add few new genes. The solid line shows the observed rarefaction curve across the 8 sampled databases, computed from 1,000 random permutations of database-addition order (seed 42); the shaded region is the 2.5th–97.5th percentile permutation envelope. The horizontal dotted line marks the Chao2 asymptotic estimate[3] of 6,865 total NDD genes, with its analytic 95% confidence interval of 6,639–7,126. At 8 databases, 5,166 unique genes are observed, corresponding to an incidence-based sample coverage[4] of 0.9116 (Supplemental Methods). Data: SysNDD API v0.38.1 (accessed September 21, 2026).

### Figure S5 | Sensitivity of Gene Count Estimates

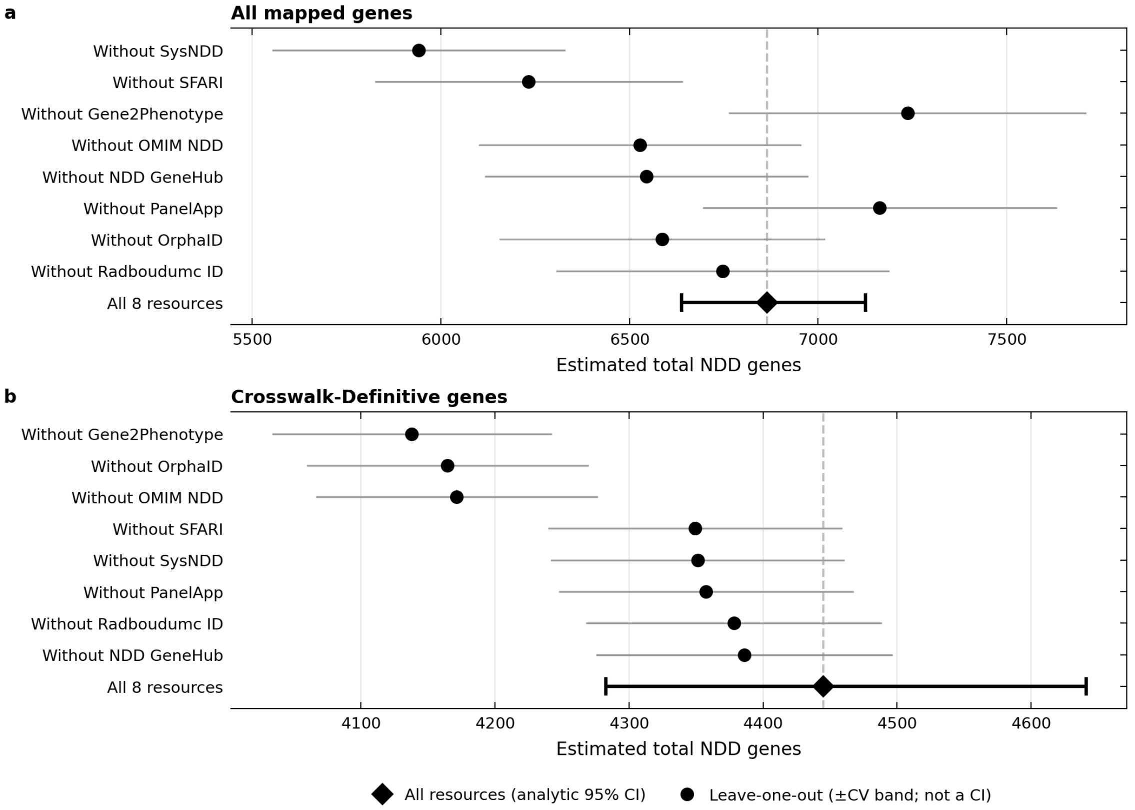

**Figure S5. Leave-one-database-out sensitivity of the Chao2 total NDD gene estimate.** Forest plots of Chao2 projections[3] of the total NDD gene count (note the different x-axis scales) for (**a**) all mapped genes and (**b**) the crosswalk-Definitive stratum (Supplemental Methods, Gene Count Estimation and Database Comparison). Rows show the projection after removing each of the eight resources in turn (black circles), ordered by the size of the change from the all-resource projection (black diamond, last row; vertical dashed line). Removing a resource lowers the projection when it lists many genes found nowhere else and raises it when it mainly repeats genes listed elsewhere. (**a**) The 5,166 observed genes project to 6,865; the leave-one-out projections range from 5,941 to 7,237 (coefficient of variation 0.0655), and removing SysNDD, the resource with the most genes listed nowhere else, lowers the projection most. (**b**) The 3,480 crosswalk-Definitive genes project to 4,445; the leave-one-out projections range from 4,138 to 4,386 (coefficient of variation 0.0252). Every removal lowers the projection, with the largest decreases after removing Gene2Phenotype, OrphaID or OMIM NDD, which contribute most of this stratum's unique genes, whereas removing SysNDD changes the projection little. The lower values in (**b**) reflect the smaller gene set rather than a different estimate of the same quantity. Error bars: analytic log-normal 95% confidence intervals for the all-resource projections (capped bars). For the other rows, capless grey bars extend on either side of the row's own projection by that projection times the coefficient of variation of the panel's leave-one-out projections, a guide to spread rather than a confidence interval. Data: SysNDD API v0.38.1 (accessed September 21, 2026).

### Figure S6 | Curation Coverage Analysis

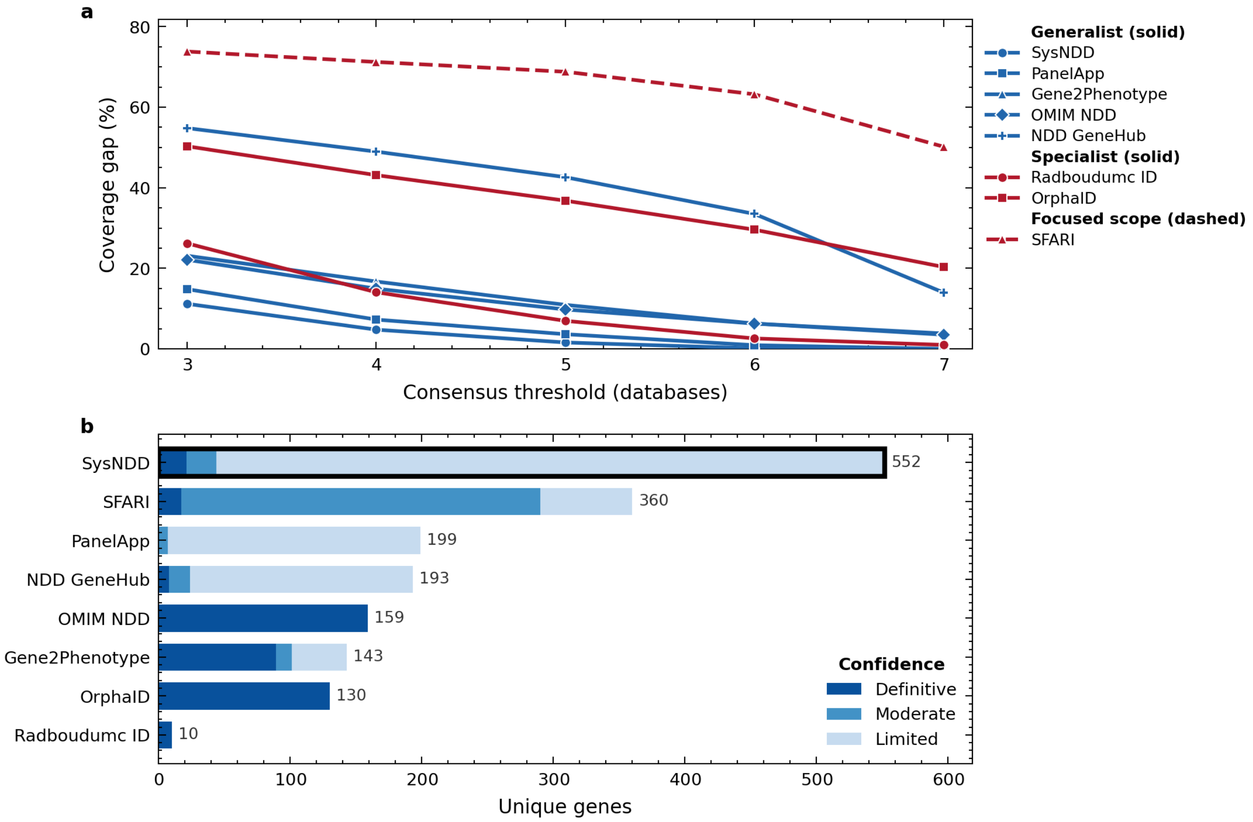

**Figure S6. Leave-one-out curation coverage analysis across eight NDD gene databases.** (**a**) Consensus coverage gap of each database at increasing agreement thresholds (3–7 databases). The gap is the proportion of genes present in at least the threshold number of the other seven databases that the database in question lacks (Supplemental Methods); lower curves indicate more complete coverage. Color marks database family (generalist or specialist) and marker shape identifies each database; line style separates the broad-scope resources (solid) from the autism-focused SFARI (dashed), whose higher gap reflects its narrower disease scope. (**b**) Genes present in only one of the eight databases, per database, stacked by crosswalk tier (crosswalk-Definitive, Moderate, Limited; Refuted assertions do not count as presence, so each bar sums to the database's unique-gene total). A bold outline marks SysNDD. Data: SysNDD API v0.38.1 (accessed September 21, 2026).

#### Figure S7 | Curation-Tier Concordance with External Standards

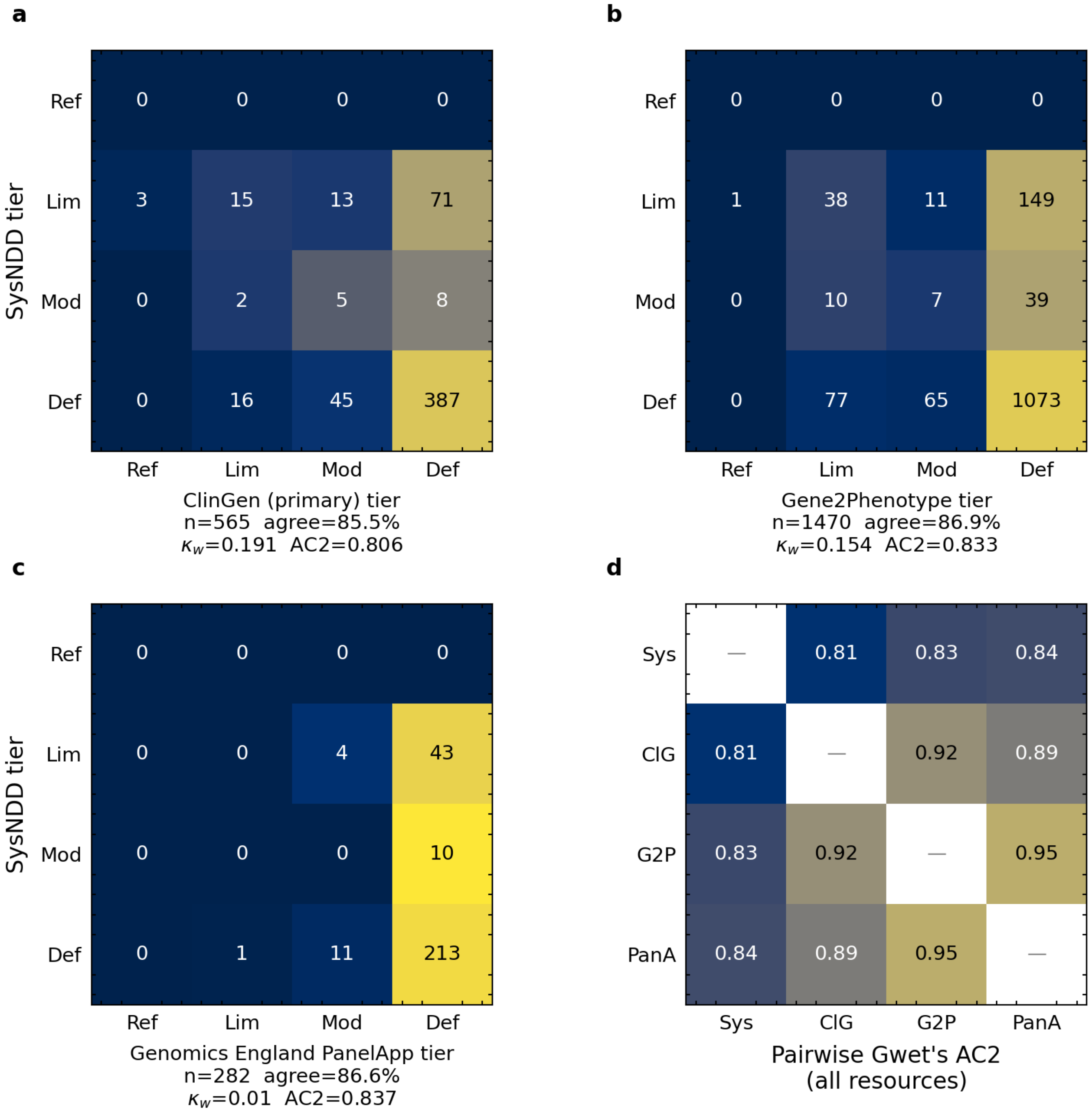

Rows: SysNDD tier; columns: comparator tier; cells: count (row-normalized shading).

**Figure S7. Concordance of SysNDD evidence categories with external gene-disease validity standards.** Confusion matrices for gene-disease pairs co-curated by SysNDD (rows) and each external standard (columns), with all resources collapsed to the SysNDD four-tier scale (Table S1). (**a**) ClinGen (the primary comparator; n = 565 pairs), (**b**) Gene2Phenotype (n = 1,470), and (**c**) Genomics England PanelApp (n = 282). Cells are shaded by row-normalized proportion (external tier given SysNDD tier), so the dominant Definitive/Definitive cell does not mask off-diagonal structure; annotations are raw pair counts. Below each panel: number of co-curated pairs, observed linear-weighted agreement, the linear-weighted Cohen's $\kappa$, and the prevalence-robust Gwet's AC2. Low $\kappa$ beside high agreement and AC2 reflects that most pairs are Definitive in both resources, which depresses $\kappa$ (Supplemental Methods). Off-diagonal mass is concentrated where SysNDD assigns the lower tier. (**d**) All-vs-all pairwise Gwet's AC2 between every resource (each pair on its own co-curated set; the diagonal is omitted). ClinGen, Gene2Phenotype, and PanelApp, which all express their assertions in GenCC's validity terms, agree most closely with one another (AC2 0.89–0.95); SysNDD is slightly less concordant with each (AC2 0.81–0.84). Data: GenCC export accessed June 21, 2026; SysNDD data release accessed September 21, 2026.

### Figure S8 | Complete Phenotype and Variant Distributions

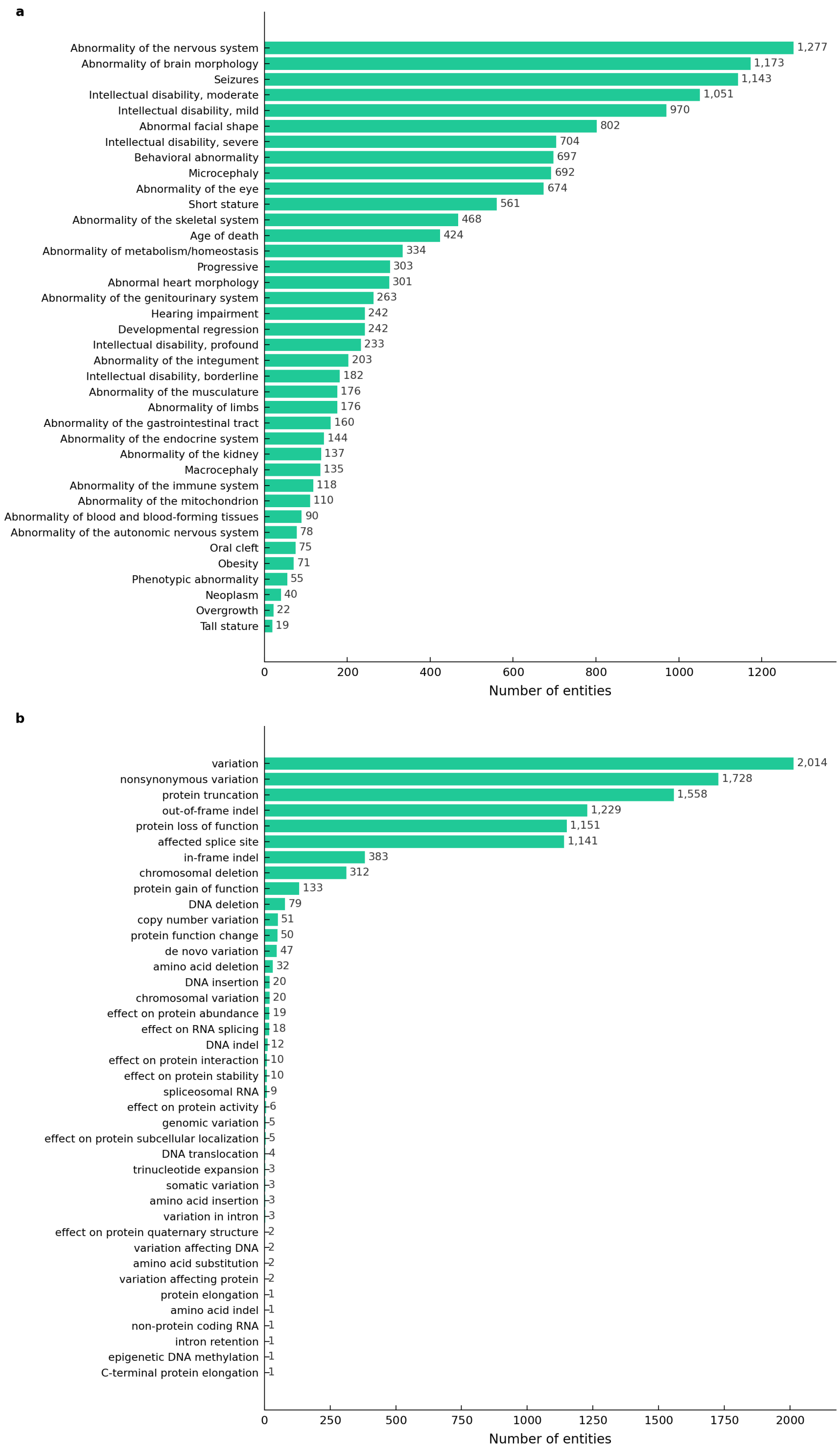

**Figure S8. Complete phenotype and variant type distributions in SysNDD.** (**a**) All 38 Human Phenotype Ontology (HPO) phenotype categories[5,6] after the base term "Intellectual disability" (HP:0001249) is excluded; the five severity grades are retained (14,545 annotations). Categories are ordered by entity count; main Figure 4a shows the top 20. (**b**) All 40 Variation Ontology (VariO) variant categories[7] (10,072 annotations). Because both ontologies are hierarchical, categories are not mutually exclusive, and one entity can contribute to several. Data: SysNDD API v0.38.1 (accessed September 21, 2026).

### Figure S9 | Recorded Syndromicity of Definitive NDD Entities

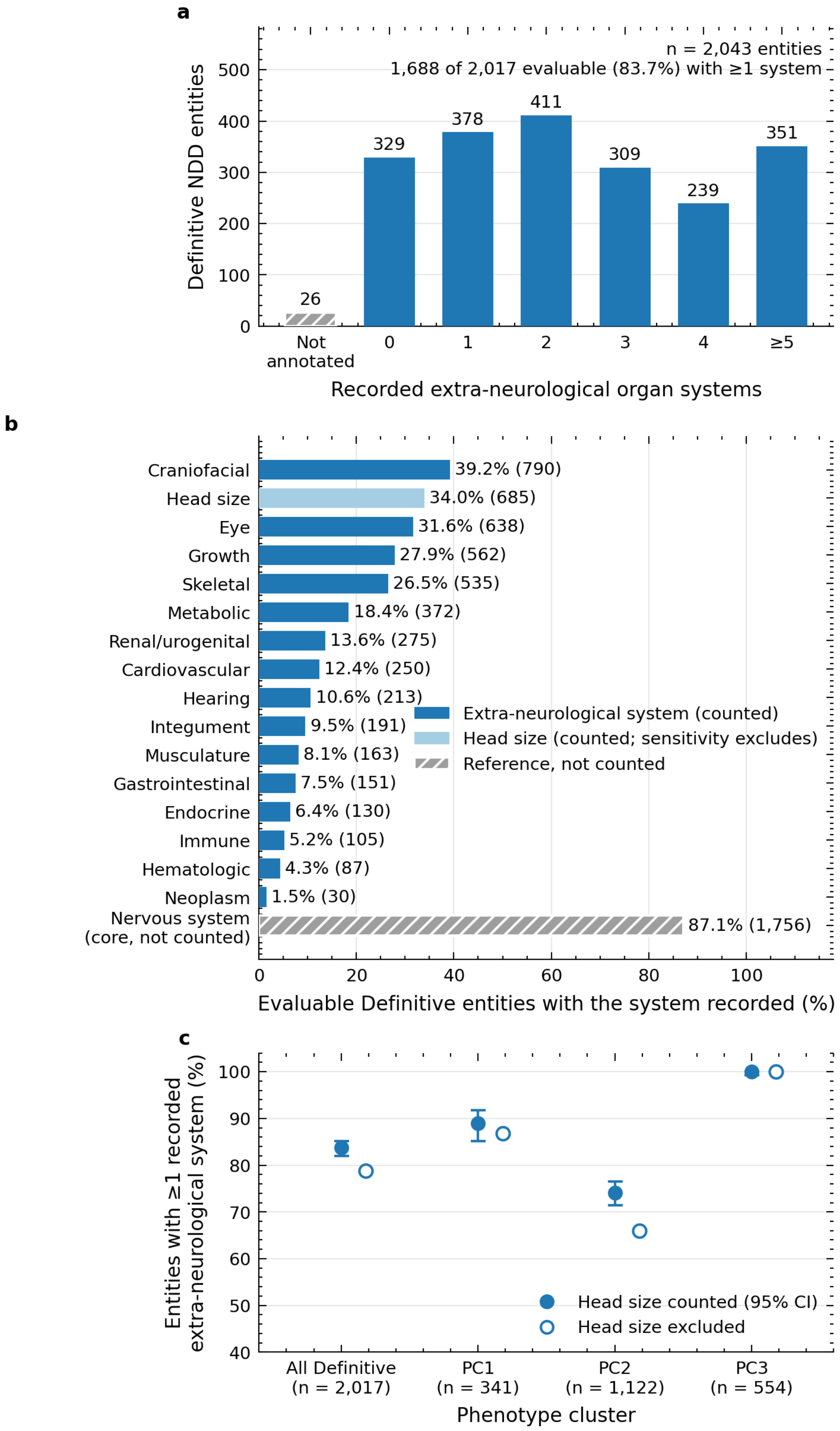

**Figure S9. Recorded syndromicity of Definitive NDD entities.** Syndromicity is recorded as extra-neurological organ involvement: 24 of the annotated phenotype terms describe such involvement and were collapsed into 16 organ systems, each counted once (Supplemental Methods). (**a**) Distribution of the 2,043 Definitive NDD entities by the number of organ systems with a phenotype recorded as present. The 26 entities without any present annotation are shown separately (hatched); the 329 entities in the zero bin carry annotations but none outside the nervous system. (**b**) Fraction of the 2,017 annotated entities with each organ system recorded. Abnormal head size is counted as an extra-neurological finding (light blue); without it, the share of entities with any recorded system falls from 83.7% to 78.8%. Nervous-system involvement (hatched grey) is the defining phenotype and is shown for reference only. (**c**) Fraction of entities with at least one recorded extra-neurological system, for all Definitive entities and for each phenotype cluster of Figure 5b. Filled markers carry Wilson 95% confidence intervals[8], which express the precision afforded by group size; open markers exclude head size. PC3 is uniformly syndromic (100% of entities), whereas recorded extra-neurological involvement is least frequent in PC2, the intellectual disability and seizures cluster (74.1%; 95% CI 71.4–76.5%), where excluding head size lowers it most (to 66%). Data: SysNDD API v0.38.1 (accessed September 21, 2026).

### Figure S10 | Full Phenotype Correlation Matrix

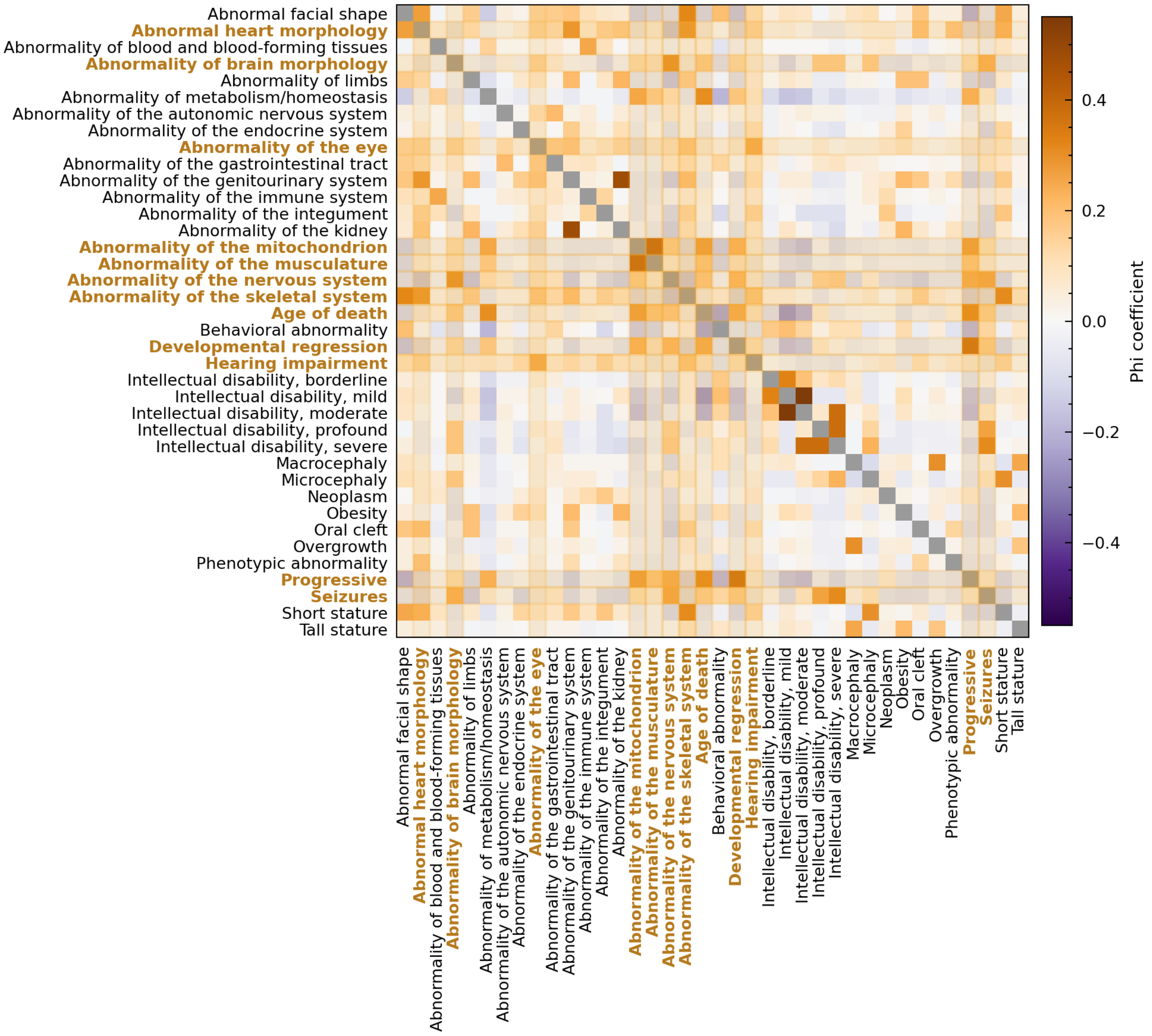

**Figure S10. Complete phenotype-phenotype correlation matrix.** Pairwise phi coefficients (Pearson *r* for binary phenotype-category presence across entities) between all 38 HPO phenotype categories annotated in SysNDD. Purple indicates negative association and orange indicates positive association; a negative coefficient denotes less frequent co-annotation than expected from the marginal frequencies. Color intensity is scaled to the observed off-diagonal range; the diagonal (each category with itself, *r* = 1) is masked in gray. Pale orange rows and columns, with the category names in bold orange type, mark the 12 phenotypes shown in the focused subset in Figure 4c. Because HPO is a hierarchical ontology, nested ancestor/descendant categories are not statistically independent. Among categories annotated in at least 10 entities, the strongest positive correlation is between Intellectual disability, moderate / Intellectual disability, mild (r = 0.55) and the strongest negative correlation between Age of death / Intellectual disability, mild (r = -0.26). Data: SysNDD API v0.38.1 (accessed September 21, 2026).

### Figure S11 | Full Variant-Consequence Correlation Matrix

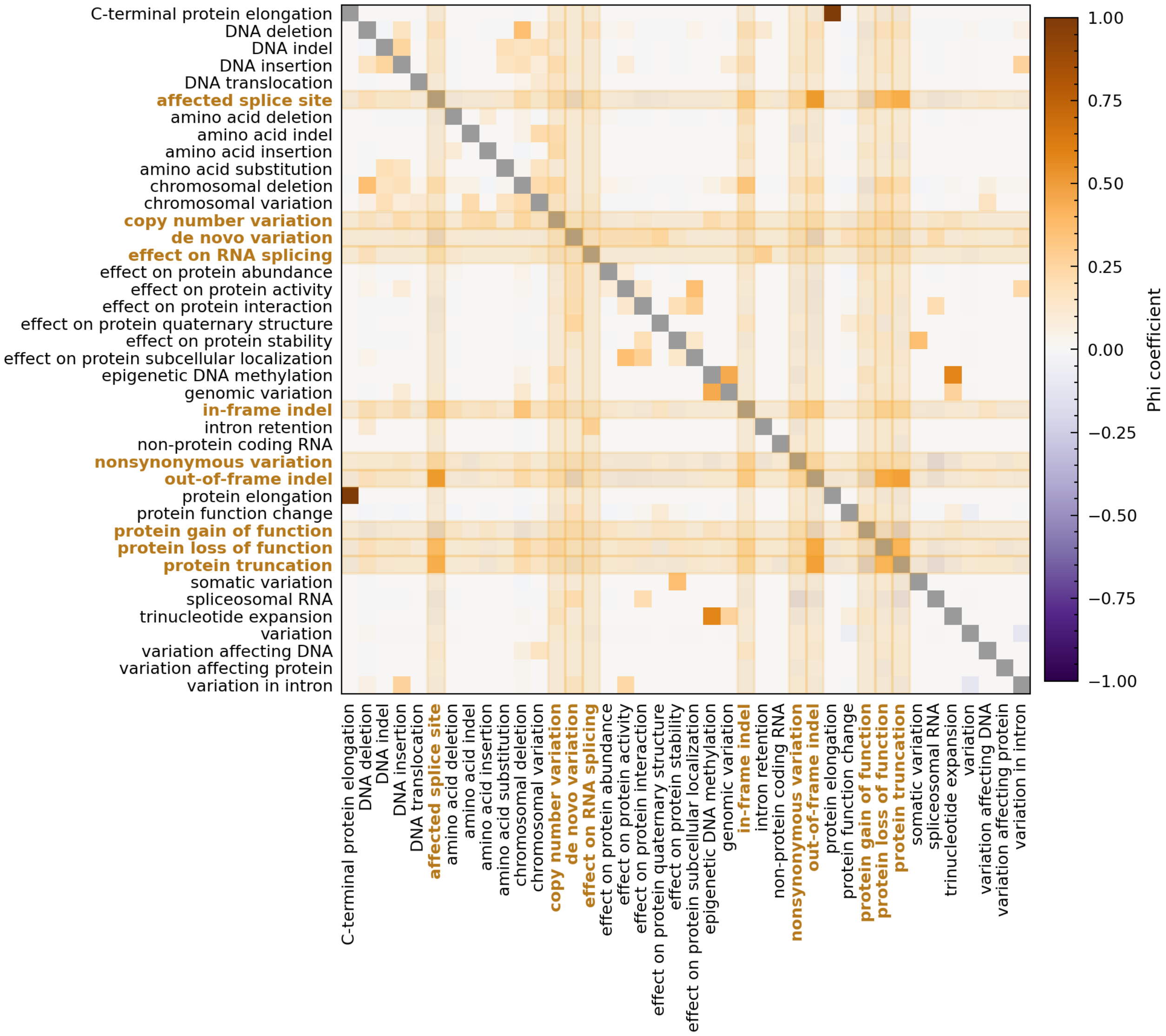

**Figure S11. Complete variant-consequence correlation matrix.** Pairwise phi coefficients (Pearson *r* for binary class presence across entities) between all 40 Variation Ontology (VariO) variant-consequence classes[7] annotated in SysNDD. Purple indicates negative association and orange indicates positive association; scaling, the diagonal masked in gray, and the reading of negative coefficients are as in Figure S10, and because VariO is hierarchical, nested classes are not statistically independent. Pale orange rows and columns, with the class names in bold orange type, mark the 10 mechanistically interpretable classes shown in Figure 4d. Among classes annotated in at least 10 entities, the strongest positive correlation is between affected splice site / out-of-frame indel (r = 0.52) and the strongest negative correlation between protein gain of function / protein truncation (r = -0.19). Some sparser classes reach larger coefficients, but are annotated in too few entities to be reliable. Data: SysNDD API v0.38.1 (accessed September 21, 2026).

### Figure S12 | Robustness of Functional-Network Modularity to STRING Channel Choice

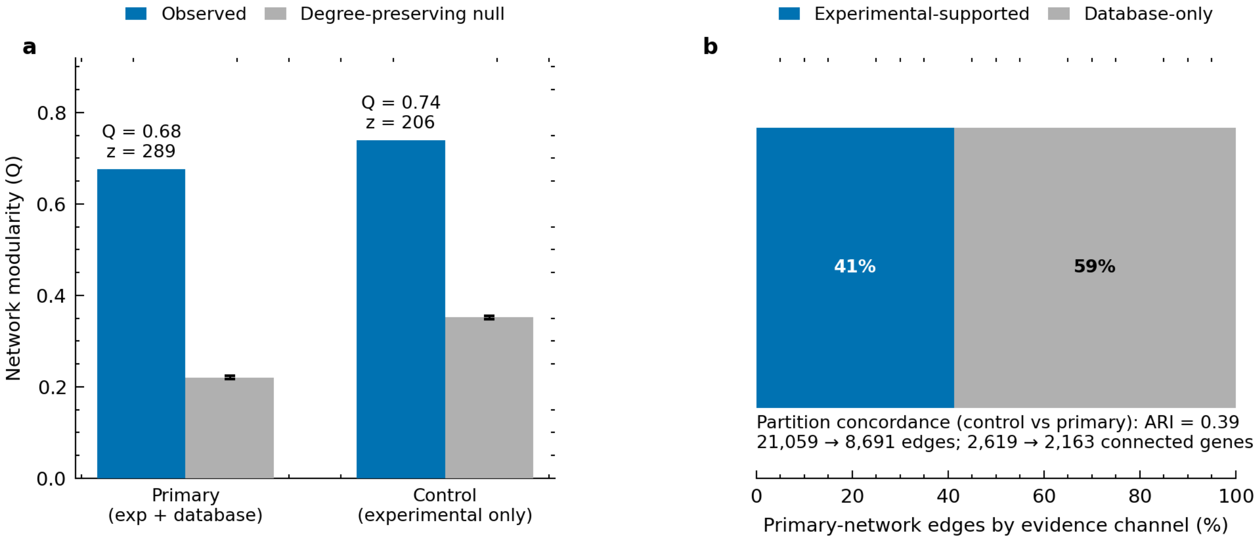

**Figure S12. Experimental-channel-only control for functional-network modularity.** The control removes STRING's curated database channel, whose pathway and complex co-membership partly overlaps the curated phenotype annotations (Supplemental Methods, Functional Clustering). (**a**) Modularity (Q) of the primary (experimental plus database) and experimental-only partitions, each against a degree-preserving, re-optimized configuration-model null[9,10] (200 replicates; grey bars, null mean; error bars, $\pm 2$ SD; z-scores annotated). The null consists of randomized networks in which every gene keeps its number of interactions. Both observed values lie far above their nulls. (**b**) Provenance of the primary network's 21,059 edges by STRING channel: 41% have experimental support and 59% rest on the database channel alone. Removing that channel leaves 8,691 edges among 2,163 connected genes; the resulting partition agrees only moderately with the primary one (adjusted Rand index[11] 0.39). Network construction and Leiden parameters are as in Supplemental Methods.

### Figure S13 | Functional Clusters against Individual Phenotype Categories

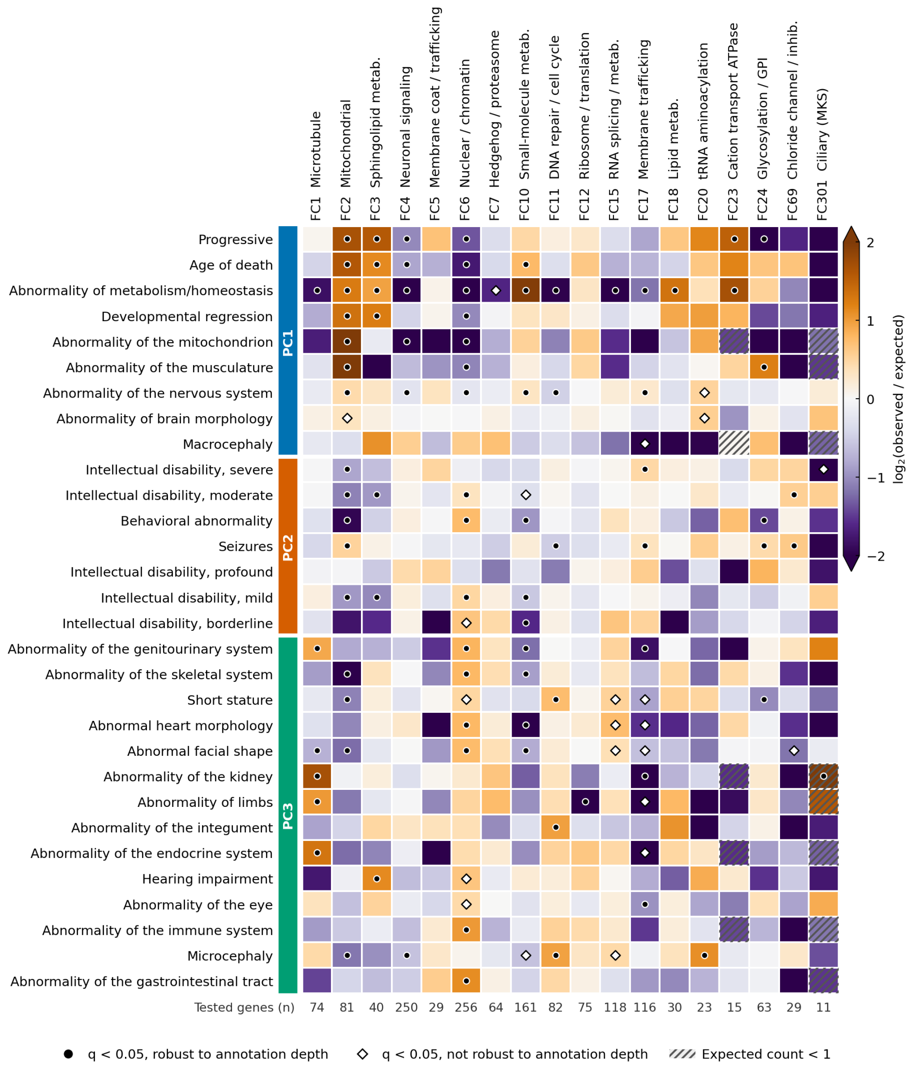

**Figure S13. Association of the functional clusters with individual phenotype categories.** Each of the 18 functional clusters (columns, identified and named as in Figure 5a) was tested against each of the 30 active phenotype categories (rows) over the 1,517 genes of the shared gene set (Supplemental Methods). The number of tested genes is given below each column. Color is the $\log_2$ ratio of observed to expected gene counts, truncated at the ends of the scale: orange indicates more genes than expected and purple fewer. Rows are grouped by the phenotype cluster of Figure 5b in which the category is most over-represented (side bar). Of the 540 cells, 110 are significant at $q < 0.05$ (two-sided Fisher exact test, one Benjamini-Hochberg correction over all cells; 55 enrichments and 55 depletions). Filled circles mark the 88 of them that remain significant when cluster labels are permuted within strata of annotation depth; open diamonds mark those that do not. Hatching marks the 13 cells with an expected count below 1, which include the enrichment of kidney abnormalities in the ciliary cluster FC301 (6 of 11 genes). A category carried by no gene of a cluster appears at the purple end of the scale whatever the size of the cluster; only marked cells are significant. The intellectual disability severity grades are not mutually exclusive, and nested categories are not independent, so the cells are not independent findings. Data: SysNDD API v0.38.1 (accessed September 21, 2026).

### Supplemental Methods

#### Phenotype Clustering

Phenotype clustering asks which Definitive entities (gene-inheritance-disease associations) resemble one another in their recorded phenotypes. It covered the 2,017 Definitive entities with at least one Human Phenotype Ontology (HPO) phenotype annotation[5,6], spanning 38 HPO phenotype categories. Entities were placed in a low-dimensional space by multiple correspondence analysis (MCA; FactoMineR R package[12]), a principal-components-like method for categorical data, and then grouped by hierarchical clustering on principal components (HCPC)[13]. The first 8 MCA dimensions were retained and a Ward tree was built on them. The number of clusters (k = 3) was not fixed in advance. It is the value of $k$ at which the step from $k-1$ to $k$ clusters gave the largest proportional drop in within-cluster inertia, the spread of entities around their cluster centers. The cut was then consolidated (refined by k-means over all entities), started from the centroids of the cut and from 100 seeded random starts (seed 42), and the converged solution with the lowest within-cluster inertia was kept.

Active variables were prevalence-filtered before MCA: categories outside a 5–95% prevalence band were moved out of the active set, leaving 30 of the 38 categories active. All removed categories fell below the 5% floor (among them the root term “Phenotypic abnormality”, neoplasm, overgrowth, and tall stature); none of the 38 categories approached the 95% ceiling. The only input removed by the ceiling was the base term “Intellectual disability” (HP:0001249), which SysNDD records in addition to these categories and which nearly all Definitive entities carry. Cognitive impairment therefore enters the analysis through the five intellectual disability severity grades, all of which remained active. In the primary analysis a category not recorded for an entity counted as absent. When unrecorded categories were instead treated as unknown (a positive-only analysis), the partition (the assignment of every entity to one cluster) was not reproduced. The adjusted Rand index[11], where 1 means identical clusters and 0 means agreement no better than chance, was near zero.

Two further analyses varied the active set. The procedure was first reproduced independently from the entity annotations: the recomputed MCA coordinates agreed with the released coordinates to within 0.00005, and the recomputed partition was identical to the released one (adjusted Rand index 1.00). Removing the five intellectual disability severity grades (25 active categories), so that only accompanying phenotypes were clustered, or omitting prevalence filtering (38 categories) again selected 3 clusters and largely retained the partition (adjusted Rand index 0.75 and 0.83 against the released partition). Neither variant raised the mean silhouette above 0.222 at any k tested. The consolidation has a second, poorer solution on these coordinates (within-cluster inertia 0.360 against 0.353; mean silhouette 0.145 against 0.183). It is reached from 20 of the 101 starts, among them the start from the hierarchical cut, which is why several starts are searched. This second solution contains the same progressive/metabolic cluster PC1 but divides the remaining entities by intellectual disability severity and seizures rather than by malformation and growth features (adjusted Rand index 0.35 against the released partition). Both reruns agree with it far less (0.25 and 0.32). The boundary between PC2 and PC3 is therefore the least certain part of the partition, as expected of a continuum, whereas the 4 strongest phenotype-functional enrichments fall in the same functional clusters under either solution.

The mean silhouette[14] was low (0.183), although well above the value obtained when cluster labels are shuffled at random (silhouette z = 49.2). The silhouette runs from $-1$ to 1: values near 0 mean that clusters overlap, negative values mean that an entity sits closer to another cluster than to its own, and 1 would mean perfect separation. The partition is therefore reproducible under resampling (bootstrap-Jaccard 0.935–0.947, where 1 means the same clusters on every resample; Table S6) but only weakly separated. Its clusters are not interpreted as discrete phenotype groups in this representation (dip tests under Cluster Structure and Separation).

#### Functional Clustering

Functional clustering asks which NDD-associated genes act together in the cell. It was based exclusively on STRING v11.5 functional-association edges[15] for all 2,601 clustered genes, irrespective of evidence category. Gene Ontology (GO)[16,17], Reactome[18], and HPO term enrichment were used only to interpret and label the resulting clusters. STRING scores each gene pair on several evidence channels. The text-mining channel was excluded, the experimental and database channels were recombined with STRING's probabilistic-OR formula (prior 0.041), and edges with a recombined score of at least 400 were kept. Weighted Leiden community detection[19] then optimized modularity[20], the share of interaction weight that falls within clusters beyond what chance would give, to convergence at resolution 1 (seed 42). Of the resulting partition, the 18 communities with at least 10 genes are reported as functional clusters; modularity values refer to the full partition.

To ask whether this modularity exceeds what the network's connectivity alone would produce, communities were re-detected in each of 200 randomized networks with the same connectivity (every gene keeps its number of interactions; a degree-preserving configuration-model null[9,10]). The observed full-partition modularity (0.679) was 259.4 standard deviations above the null mean (0.22; empirical $p <= 0.005$). Modularity of the largest connected component, which also retains genes in communities below the 10-gene threshold, was 0.678.

Excluding text mining leaves 550 genes with only literature co-mention evidence as isolates and reduces co-study bias, but STRING's database channel still overlaps clinical curation. An experimental-channel-only control (Figure S12) therefore rebuilt the network from the STRING v11.5 detailed-links file without that channel, restricted to the largest connected component, with the same prior, threshold, resolution and seed. Modularity remained far above its degree-preserving null (Q = 0.74, null mean 0.35, z = 206), but the experimental-only partition agreed only moderately with the primary one (adjusted Rand index 0.39). The modular organization as a whole is therefore not an artifact of the curated channel, whereas individual cluster memberships depend on the channel.

To ask whether the four couplings highlighted in Figure 5c persist in that experimental-only reconstruction, each highlighted functional cluster was mapped to the experimental-only community that captures the largest share of its genes (the share given below). The corresponding phenotype cluster was then retested against that community over the shared gene set of this reconstruction, 1,205 genes (6 of 81 phenotype-cluster-by-community cells significantly enriched at $q < 0.05$). Of the four, 3 persist: PC1–FC2 (dominant-community share 70%, OR = 21.804, $q < 0.001$), PC3–FC6 (share 68%, OR = 2.351, $q < 0.001$), and PC2–FC17 (share 50%, OR = 2.087, $q = 0.018$). PC1–FC10 does not (share 35%, OR = 1.818, $q = 0.166$).

Cluster stability was assessed by subsampling without replacement (100 resamples at 80% of entities or genes, seed 42) with cluster-wise maximum-Jaccard recovery[21]; following that method, these subsampling stability values are called bootstrap-Jaccard throughout. The resulting per-cluster bootstrap-Jaccard means, HCPC silhouettes, and Leiden partition modularity are reported in Tables S6 and S7 and Figure 5.

Stability is cluster-specific: 9 of the 18 clusters fall below bootstrap-Jaccard 0.6, and the smallest cluster (FC301, 16 genes) has bootstrap-Jaccard 0.627 (per-cluster values in Tables S6 and S7).

The two partitions, their validation metrics, and the phenotype-functional associations all derive from the same dated public data release. The Leiden partition was computed by SysNDD application release v0.37.0, the phenotype partition and the associations by release v0.38.1; all are served by API v0.38.1, which provides the largest-component edge list, both cluster memberships, and the phenotype-axis MCA coordinates.

#### AI-Generated Pathway Summaries

A natural-language pathway summary of each functional cluster was generated with the large language model Google Gemini (3.5 Flash) from the cluster's members, GO term enrichment, and network topology, and is displayed in the SysNDD Gene Networks view (https://sysndd.dbmr.unibe.ch/Analyses/GeneNetworks). The analysis release dated September 21, 2026, contains 18 such summaries, each labeled with the model version and generation date. Each passed an automated model-based check before release; the summaries are not curator-reviewed.

#### AI-Generated Phenotype Cluster Summaries

A summary of each phenotype cluster was generated with the same model from the cluster's phenotype categories and inheritance composition, and was released, checked, and labeled in the same way. The release contains 3 phenotype-cluster summaries (21 summaries in total).

#### NDDScore Prediction Layer

NDDScore is a separately developed exploratory resource that uses SysNDD evidence categories as training labels. The active release (nddscore_20260517_public, May 17, 2026) scores 19,296 protein-coding genes and predicts inheritance and HPO annotations; access and archive are listed in Table S3. Detailed NDDScore development and performance evaluation are outside the scope of this manuscript.

#### Database Comparison

SysNDD gene content was compared with seven other neurodevelopmental disorder gene resources (Results): PanelApp[22], Gene2Phenotype[23], OMIM NDD[24], OrphaID[25], SFARI[26], Radboudumc ID, and Geisinger NDD GeneHub[27] (sources and access dates in Table S9). Records were mapped to HGNC symbols[28] and harmonized by the versioned tier crosswalk (Table S1) into a single gene-by-resource table (Data S1, sheet "Gene by resource"). OMIM NDD was derived from OMIM genemap2 filtered through the HPO phenotype-to-genes table[5], in which HP:0012759 annotations are propagated to descendant phenotypes.

Because each resource grades evidence on its own scale, the native labels are harmonized to the SysNDD four-tier scale (Definitive, Moderate, Limited, Refuted), and genes absent from a resource are recorded as not listed. PanelApp Green, Amber, and Red map to Definitive, Moderate, and Limited; no PanelApp tier maps to Refuted; only source-native disputed or refuted assertions (as in Gene2Phenotype) do. SFARI scores 1–3 and Geisinger NDD GeneHub tiers map onto the same scale; the ungraded inclusion lists (Radboudumc ID, OMIM NDD, OrphaID) are carried as Definitive. Categories mapped to Definitive under this crosswalk, including the ungraded lists, are termed crosswalk-Definitive, as distinct from source-native Definitive assertions. The versioned crosswalk is published with the comparison data at https://sysndd.dbmr.unibe.ch/CurationComparisons.

Pairwise cosine similarities (1 when two resources list exactly the same genes, 0 when they share none) were calculated from binary gene-membership vectors with the R coop package[29]. The similarities were converted to cosine distances ($1 -$ similarity), and average-linkage hierarchical clustering was performed with SciPy[30]. Multi-database intersections were visualized as UpSet plots[2].

#### Technical Implementation

SysNDD consists of a Vue.js web interface, an R plumber API[31] documented with an OpenAPI specification, and a MySQL database; the application is deployed as Docker containers.

#### Read-only Model Context Protocol Interface

The Model Context Protocol interface gives a language-model assistant read-only access to the same public records and analysis releases as the REST interface, so that it can retrieve and cite curated evidence by record identifier. The records exposed are entities with their evidence and inheritance, supporting publications as PMIDs, and analysis releases. It excludes draft records and administrator functions and does not generate text. It is not a separate evidence source and was not evaluated as a clinical decision-support system.

#### Web Interface

The public web application provides gene-, disease-, phenotype-, and entity-centered access to approved SysNDD records. Tabular views for entities, genes, phenotypes, and panels support searching, filtering, sorting, and export; the Phenotypes view combines HPO categories with AND/OR logic, and the Panels view builds downloadable gene lists by evidence category and inheritance.

Public analysis views cover database comparison, phenotype and variant annotation patterns, temporal growth, curated-publication statistics, PubTator literature annotations[32], and the functional and phenotype clusterings, with downloadable analysis releases. These views label analysis outputs as derived from a dated SysNDD release, distinct from approved curation.

Authenticated access requires registration and approval. A Reviewer proposes changes to assigned re-review records (synopsis, phenotype and variation annotations, literature links, and evidence category) or flags an entity for specialist attention. A submission does not itself replace the public active record until a Curator or Administrator approves it. Curator and Administrator roles create or modify entities, approve pending reviews and statuses, evaluate account applications, allocate re-review work, and maintain GeneReviews coverage. Administrator pages maintain user access, vocabularies, external annotations, explanatory content, derived and NDDScore prediction releases, operational records, and database recovery, all separated from evidence-category assignment. Changes that could affect curated evidence are staged, retained for re-review, validated before activation, or published as immutable versions. Table S2 lists every page family with its access role, purpose, and data-control boundary.

Curation criteria and full usage documentation are available at https://berntpopp.github.io/sysndd/05-curation-criteria.html.

#### Statistical Analysis

##### Annotation Co-occurrence

Co-occurrence of annotation categories (Figure 4c and 4d; Figures S10 and S11) was computed over Definitive NDD entities with at least one recorded annotation of the relevant type, each encoded by the presence or absence of every phenotype category (a category not recorded for an entity counts as absent) or every VariO class; the base term "Intellectual disability" (HP:0001249) was excluded in favor of its severity grades. For each pair of categories, the phi coefficient (the Pearson correlation of the two binary vectors) was calculated. A positive value means that two categories are recorded together more often than their individual frequencies predict, and a negative value that they are recorded together less often. Phi can reach 1 only for two categories of equal frequency. For categories with prevalences $p_1 \leq p_2$ its attainable positive maximum is $\sqrt{p_1(1-p_2)/[p_2(1-p_1)]}$, so a moderate value between a rare and a common category can lie close to the strongest association possible. The coefficients are reported descriptively, without significance testing or correction for multiple comparisons, and because both ontologies are hierarchical, nested categories are not independent.

##### Phenotype-Functional Cluster Association Testing

These tests ask whether the genes of a phenotype cluster are over- or under-represented in a functional cluster. Genes were the unit of analysis: phenotype clusters are defined over entities and functional clusters over genes, so entity membership was collapsed and each gene counted once per phenotype cluster. The tests were restricted to the shared gene set, the 1,517 genes present in both clusterings, out of 1,823 genes in the phenotype clusters and 2,601 in the functional clusters.

Each of the 54 phenotype-by-functional cells was summarized by its 2 Œ 2 table (a, genes in both clusters; b, phenotype cluster only; c, functional cluster only; d, neither). From that table we report the Fisher exact odds ratio, which measures how strongly the two memberships coincide (1 means independence), with its log-scale 95% confidence interval. We also report the Jaccard overlap (the shared genes as a fraction of the genes in either cluster), the Fisher exact p-value, and a seeded size-preserving permutation p-value.

When any cell was zero, the odds-ratio confidence interval used the Haldane-Anscombe correction[33,34].

The two-sided permutation null used 10,000 replicates with seed 42. Each replicate redrew every phenotype cluster as a random gene set of the same size from the shared gene set, holding the functional clusters constant. The permutation p-value is the share of replicates whose overlap lay at least as far from the null expectation as the observed overlap, with the observed overlap counted among them.

Benjamini-Hochberg correction[35], which limits the false discovery rate (the expected share of false positives among the cells called significant), was applied once over the full family of 54 PC–FC cells using the Fisher exact p-values, at $q < 0.05$. The permutation test serves only as a robustness check. Pearson coefficients appear only as heatmap colors in Figure 5c and were not used for inference. The complete per-cell table (2 Œ 2 counts, odds ratios with 95% confidence intervals, Jaccard overlaps, Fisher exact and permutation p-values, q-values, and direction) is provided in Data S1 (sheet "PC-FC associations").

Restricting cluster membership to Definitive genes (Table S4) changed neither the shared gene set nor any 2 Œ 2 table, because only genes present in both clusterings are tested and every phenotype-cluster gene is already Definitive. The functional network was not re-clustered, so this check does not show that the functional partition itself is independent of evidence category.

Because this comparison inherits a weakly separated phenotype partition, each functional cluster was also tested directly against the 30 active phenotype categories (Figure S13). Genes were again the unit of analysis, over the same shared gene set, and a gene carried a category when any of its Definitive entities recorded it as present. Each of the 540 cluster-by-category cells was tested with a two-sided Fisher exact test, with one Benjamini-Hochberg correction over all cells ($q < 0.05$); enrichments and depletions are both reported. A gene with more annotated categories is more likely to carry any one of them, and annotation depth differs between clusters. A permutation null (10,000 replicates, seed 42) therefore reassigned cluster labels only among genes with a similar number of active categories (5 strata). A cell was counted as robust to annotation depth when it stayed significant, in the same direction, after the same correction of the permutation p-values (88 of 110 cells). The complete per-cell table is provided in Data S1 (sheet "FC-phenotype associations").

#### Cluster Structure and Separation

Cluster structure was assessed by biological interpretability and by per-cluster stability, Leiden partition modularity[20], the HCPC mean silhouette[14] over the MCA coordinates, and a silhouette-versus-k curve, all for the same versioned data release (Figure 5; Tables S6 and S7).

The phenotype silhouette-versus-k curve uses the same consolidated HCPC procedure as the reported partition, so the curve value at the selected k equals the reported mean silhouette. Across all k tested ($k = 2$–$10$) the mean silhouette remains low ($\leq 0.18$), so the weak separation does not depend on the choice of k within this representation.

Separation is also reported within each axis as a z-score, the number of standard deviations by which an observed value exceeds the mean of its null model. For the functional partition this is the modularity z; for the phenotype partition it is the silhouette z together with a modularity z on a k-nearest-neighbour graph of the MCA coordinates (95.8). Because these z-scores use different null models on different graphs, they are not directly comparable across axes; the contrast between strong functional modularity and weak phenotype separation rests on these within-axis calibrations. As an additional check on both distance distributions, Hartigan dip tests[36], which ask whether a distribution has more than one peak, were run on the MCA-coordinate distances and on the functional interaction distances. Neither test rejected unimodality ($p = 0.946$ and $p = 0.3902$). These results are not evidence of significant multimodality; because the tests run on mutually dependent pairwise distances, they are descriptive checks rather than calibrated evidence for discrete biological classes.

##### Recorded Syndromicity

Syndromicity (Figure S9) was defined as recorded extra-neurological organ involvement and computed over all Definitive NDD entities: every Definitive entity whose disease is recorded as neurodevelopmental, each counted once. SysNDD annotates phenotypes against a fixed vocabulary of 38 categories plus the base term “Intellectual disability” (HP:0001249), and each of these 39 terms was assigned one role (Data S1, sheet “Syndromicity registry”). Terms that do not describe additional organ involvement were excluded: the ontology root “Phenotypic abnormality” (HP:0000118), the clinical-course terms “Progressive” (HP:0003676) and “Age of death” (HP:0011420), and the intellectual disability term with its five severity grades (borderline, HP:0006889; mild, HP:0001256; moderate, HP:0002342; severe, HP:0010864; profound, HP:0002187). The 6 nervous-system terms (abnormality of the nervous system, behavioral abnormality, seizures, abnormality of brain morphology, abnormality of the autonomic nervous system, and developmental regression) describe the defining phenotype of an NDD and were tallied separately as neurological involvement.

The remaining 24 terms were collapsed into 16 organ systems so that nested or overlapping terms count once. Six systems merge terms: renal/urogenital (kidney, genitourinary), skeletal (skeletal, limb), metabolic (metabolic, mitochondrial), growth (short stature, tall stature, overgrowth, obesity), craniofacial (oral cleft, abnormal facial shape) and head size (microcephaly, macrocephaly); eye, hearing, endocrine, integument, heart, blood, immune, musculature, gastrointestinal tract, and neoplasm each form their own system.

Only annotations recorded as present counted as evidence; annotations recorded as uncertain, variable, or rare, and the 2 annotations of explicit absence among Definitive entities, were not counted. An entity was classified as having extra-neurological involvement when at least one organ system was annotated as present. It was classified as having none recorded when it carried present annotations but no such system; the label “isolated” is avoided because this class describes documentation, not a clinical finding. Entities without any present annotation were left unclassified (per-entity classifications in Data S1, sheet “Syndromicity entities”). Proportions are reported over classified entities. Every curated entity is counted, so within the catalog these percentages are exact. The Wilson 95% confidence intervals[8] instead treat the curated entities as a sample of all Definitive NDD entities, including those yet to be curated, and show the precision that each group size affords. Because counting abnormal head size as extra-neurological is the most contestable mapping choice, every proportion is also reported with head size excluded. The per-cluster values in Figure S9c and Table S6 are published with each SysNDD phenotype clustering release under the same rule (SysNDD API v0.38.1, accessed

September 21, 2026).

##### Reporting Conventions

Exact p-values are reported where p $\geq$ 0.001 and as p < 0.001 otherwise. FDR-corrected q-values are reported to three decimal places, with values below 0.001 shown as <0.001. Correlation coefficients are reported to two decimal places with 95% confidence intervals where applicable; phenotype-functional overlap magnitudes are reported as odds ratios and Jaccard indices.

#### Gene Count Estimation

To project how many NDD genes the compared resources have yet to list, the total NDD gene count was estimated with species richness estimators from biodiversity statistics. Each of the 8 compared databases (Database Comparison) was treated as one sampling occasion of a single underlying gene population, and a gene's presence or absence in each database was treated as its incidence record. Estimates were calculated for all mapped genes and for the crosswalk-Definitive subset.

The Chao2 estimator[3], a non-parametric incidence-based richness estimator, is defined as:

$$\hat{S}_{\text{Chao2}} = S_{\text{obs}} + \frac{T-1}{T}\frac{Q_1^2}{2Q_2}$$

where $S_{\text{obs}}$ is the number of observed genes, $T$ is the number of databases, $Q_1$ is the number of genes appearing in exactly one database (incidence uniques), and $Q_2$ is the number appearing in exactly two databases (incidence duplicates). When $Q_2 = 0$, the correction term is replaced by $[(T-1)/T]Q_1(Q_1-1)/2$. Chao2 infers unobserved richness from the frequency of rare observations: how many genes remain unlisted is inferred from how many appear in only one or two resources. It is a lower bound only when the sampling units are independent and comparable. Overlapping curator networks and differing resource scopes violate that assumption here, so the value is reported as a descriptive projection, not as a formal lower bound or a count of undiscovered genes.

The 95% confidence interval was calculated from the analytic Chao2 variance using a log-normal transformation of the estimated number of unobserved genes[3], so that the lower bound of the interval cannot fall below the observed gene count.

Incidence-based sample coverage, the proportion of the total incidence probability represented by the observed genes, was estimated as[4]:

$$\hat{C} = 1 - \frac{Q_1}{U}\left[\frac{(T-1)Q_1}{(T-1)Q_1 + 2Q_2}\right]$$

where $U$ is the total number of gene incidences across all databases. This quantity measures incidence completeness and should not be interpreted as the proportion of all NDD-associated genes already observed.

Leave-one-database-out sensitivity analysis recalculated Chao2 after removing each database in turn, in both strata (Data S1, sheet “Chao2 estimates”).

#### Curation Coverage Analysis

Curation coverage asks which genes a database lacks although a given number of the other databases list them. For each of the 8 databases, a consensus gene set was defined as the genes appearing in at least $k$ of the remaining databases ($k$ from 3 to 7). The consensus coverage gap is the share of that consensus set missing from the database in question:

$$\text{Coverage Gap}_k = \frac{|\text{Consensus}_k \setminus \text{Database}|}{|\text{Consensus}_k|}$$

where $\text{Consensus}_k$ is the set of genes appearing in $\geq k$ databases (excluding the target), and Database is the gene set of the target database. Lower coverage gaps indicate more complete coverage of consensus NDD genes.

Unique gene contributions are genes present in exactly one database, stratified by crosswalk tier (crosswalk-Definitive, Moderate, Limited) where graded; these contributions flag database-specific curation, whether from specialized literature coverage or divergent criteria.

Both strata (all mapped genes; crosswalk-Definitive genes) were analyzed (Figure S6; Data S1, sheets “Coverage gaps” and “Unique contributions”).

The richness estimators and the coverage analysis were implemented in Python 3.11.15 with NumPy[37], SciPy[30], and pandas[38] (Table S8).

#### Curation-Tier Concordance and Discordance Adjudication

This analysis asks how far SysNDD’s evidence categories agree with the validity classifications of other resources for the same gene-disease pairs. SysNDD was compared with ClinGen[39,40] (the prespecified primary comparator), Gene2Phenotype[23], and Genomics England PanelApp[22], all taken from the GenCC bulk submissions export[41] (accessed June 21, 2026; SHA-256 prefix 7a704cca3cad), against a fixed SysNDD data release (accessed September 21, 2026). SFARI and binary inclusion-list resources assign no graded validity tier and were excluded. Tiers were collapsed to the SysNDD four-tier ordinal scale (Definitive > Moderate > Limited > Refuted; Table S1), and diseases were matched on the GenCC export’s own harmonized MONDO identifier[42] without additional mapping. These comparisons quantify agreement after an approximate crosswalk; they are not accuracy benchmarks or gold-standard validation.

For each pairing, using the shared co-curated pairs, linear- and quadratic-weighted agreement, weighted Cohen’s $\kappa$[43], and Gwet’s AC2[44,45] were computed. AC2 is reported because $\kappa$ is depressed when most pairs fall into one category, as they do here with Definitive, even when observed agreement is high (the “$\kappa$ paradox”[46,47]); AC2 is an agreement coefficient built to resist this. Confidence intervals (95%) were obtained by nonparametric bootstrap (2,000 resamples, seed 42). The same statistics were computed pairwise between all resources on each pair’s own co-curated set (Figure S7d).

Every discordant pair was then adjudicated to distinguish genuine tier disagreements from comparison artifacts. Adjudication was performed by a large language model, Anthropic Claude (Opus 4.8, Fable 5, Opus 5 and Fable 5.1), under a fixed, versioned prompt (version v1, SHA-256 prefix dbc58afed6a6) reproduced verbatim in Note S1. Each discordance was adjudicated independently, without access to the other cases. Judgments were grounded in SysNDD curated entities, NCBI GeneReviews disease descriptions, and ClinGen gene-validity records; evidence text was treated as data, not instructions. Each verdict assigned one category (genuine tier disagreement, curation-date lag, NDD-component-versus-whole-disease scope mismatch, disease-ontology granularity, identifier/inheritance mapping error, or other), a direction (which resource held the higher tier), and the evidence consulted. One verdict per discordance is retained in Data S1 (sheet “Discordance verdicts”); verdicts were not aggregated over repeated samples. Because model output is not deterministic, the verdicts were treated as a measurement and validated by blinded expert re-adjudication.

All 579 discordances across the three standards were adjudicated, a full census rather than a sample. For validation, one author (B.P.), a geneticist and curator, re-adjudicated a comparator-stratified random sample (seed 42) of 60 discordances (20 per standard, balanced on the model’s genuine-versus-artifact verdict), assigning genuine versus artifact plus a category. The re-adjudication was blind to the model’s verdict, category, and rationale. It used the same evidence the model had received (all SysNDD entities and comparator records for the gene, with their curation and classification dates) and primary sources where needed (Data S1, sheet “Validation sample”).

Expert and model agreed on 71.7% of genuine-versus-artifact calls (Cohen’s $\kappa$[48] = 0.433) and on 53.3% of category assignments ($\kappa$ = 0.333).

Disagreements ran mostly one way. The model classed 50% of the sampled discordances as genuine and the expert 35%. In 13 of the 17 discordant cases the expert reclassified the model’s genuine call as an artifact, most often (9 cases) a scope mismatch between an NDD component and the whole disease. The census proportions in Table S5c are therefore model-assisted estimates that likely overstate the genuine share, not expert re-curation.

### Supplemental Tables

#### Table S1 | SysNDD, ClinGen and GenCC Tier Crosswalk

**Table S1. Approximate crosswalk between SysNDD evidence categories and ClinGen/GenCC gene-disease validity terminology.** ClinGen and GenCC provide community terminology for gene-disease validity assessment[41,49,50]. The mapping is intended to facilitate interpretation across resources and is not a one-to-one conversion, because SysNDD curates NDD-focused gene-inheritance-disease entities and retains non-NDD entities for completeness.

| SysNDD category | SysNDD definition | Approximate ClinGen/GenCC terminology and interpretation |
|---|---|---|
| Definitive | Established NDD gene-disease entity with strong replicated or expert-confirmed evidence | ClinGen/GenCC Definitive or Strong; broadly comparable high-confidence categories. |
| Moderate | Published evidence supports the association but is less extensive than for Definitive | ClinGen/GenCC Moderate; a comparable intermediate validity tier requiring periodic re-review. |
| Limited | Published candidate association with limited or early evidence | ClinGen/GenCC Limited; candidate-level evidence, not clinical actionability. |
| Refuted | Evidence contradicts or no longer supports the asserted association | ClinGen Disputed or Refuted; GenCC Disputed Evidence or Refuted Evidence. SysNDD combines these distinctions in one Refuted category. |
| Not applicable | Non-NDD entity retained for completeness outside the NDD evidence categories | Not mapped or outside NDD scope; distinct from “No Known Disease Relationship”, which is a validity assertion for a specified disease. |

#### Table S2 | SysNDD Web Application Pages, Access Roles, and Data-Control Boundaries

**Table S2. SysNDD web application pages, access roles, and data-control boundaries.** Page families are grouped only when they share a parent view and scientific purpose. Reviewer+ denotes Reviewer, Curator, or Administrator access; Curator+ denotes Curator or Administrator access. Public pages are available without authentication. The final column distinguishes approved curation from derived analyses, external annotations, predictions, generated summaries, and operational functions.

**(a) Public and account-access pages.**

| Page or view | Access | Purpose | Data and control boundary |
|---|---|---|---|
| Home | Public | Search genes, diseases, identifiers, and entities; inspect current counts, recent entities, and the entity concept. | Counts and recent entries summarize approved curated records; the page is read-only. |
| Search results | Public | Resolve a query across genes, diseases, identifiers, and entities and open the corresponding detail page. | Search resolves existing public records and does not create or modify them. |
| Entities | Public | Browse gene-inheritance-disease entities with field filters, sorting, pagination, row expansion, links, and spreadsheet export. | Rows represent active public entity records; export preserves the applied query. |
| Genes | Public | Aggregate and browse associated entities by gene, with filters, links, and spreadsheet export. | Gene rows summarize active public entities and do not constitute a separate curation layer. |
| Phenotypes | Public | Retrieve entities by HPO category combinations and inheritance criteria using AND/OR matching. | Results derive from approved phenotype annotations; overlapping ontology categories are not independent traits. |

| Page or view | Access | Purpose | Data and control boundary |
|---|---|---|---|
| Panels | Public | Construct and export configurable gene lists by evidence category and inheritance for use in external variant analysis. | The panel is a query-derived list, not a clinical test or a new curated record. |
| Entity detail | Public | Inspect the gene, inheritance, disease, active evidence category, synopsis, phenotype and variation annotations, and supporting literature for one entity. | Displays the approved active record and its linked evidence. |
| Gene detail | Public | Inspect associated entities, identifiers, constraint data, clinical resources, model-organism information, ClinVar records, and genomic and protein visualizations. | SysNDD entities remain curated evidence; external ClinVar and reference data are displayed alongside and are not imported into those entities. |
| Disease ontology detail | Public | Inspect disease identifiers, cross-ontology links, and associated SysNDD entities. | Ontology metadata organizes public entities but does not assign their evidence tiers. |
| Curation comparisons | Public | Compare resource overlap, cosine similarity, and gene-by-resource membership under the declared confidence crosswalk. | This is a derived analysis of released resource data; mapped tiers are not new source-native assertions. |
| Phenotype correlations | Public | Explore phenotype-category correlations, counts, and phenotype clusters. | Outputs are derived analysis of approved SysNDD phenotype annotations. |

| Page or view | Access | Purpose | Data and control boundary |
|---|---|---|---|
| Variant correlations | Public | Explore variation-category correlations and counts. | Outputs are derived analysis of approved SysNDD variation annotations and do not establish causal mechanisms. |
| Unified analysis workspace | Public | Move among synchronized phenotype-cluster, gene-network, and correlation views while retaining shared filters. | The workspace combines released derived analyses without changing their source records. |
| Entries over time | Public | Aggregate curated entities or genes over time by evidence category or inheritance. | Dates describe the SysNDD curation record and are not direct dates of biological discovery. |
| NDD publications | Public | Browse SysNDD-curated publications, their temporal distribution, and publication-level source statistics. | Literature is linked to approved reviews; the view does not grade a publication independently. |
| PubTator NDD | Public | Explore externally annotated NDD publications through publication, gene, and summary-statistic views. | PubTator supplies external literature annotations; they are not approved SysNDD entity evidence. |
| Functional gene clusters | Public | Inspect released network communities, enrichment terms, gene membership, and available cluster summaries. | Network and enrichment results are derived; summaries are model-generated, labeled as such, and not curator-reviewed evidence. |
| Phenotype-functional correlation | Public | Inspect the released association matrix between phenotype-derived and functional gene clusters. | Associations are a derived analysis and do not modify either clustering or any entity. |

| Page or view | Access | Purpose | Data and control boundary |
| --- | --- | --- | --- |
| Analysis releases | Public | Download immutable analysis bundles and inspect their release manifests and file checksums. | Each release is immutable, content-addressed, and separate from the continuously updated curation database. |
| NDDScore | Public | Inspect genome-wide gene predictions, gene-level predicted annotations, and the model card. | NDDScore is a machine-learning prediction layer, not a SysNDD evidence tier, and cannot alter curated classifications. |
| REST API explorer | Public | Inspect interactive programmatic documentation and query public resources; authenticated users can access functions allowed by their role. | Anonymous results expose public data, whereas protected functions retain the same role and approval boundaries as the web interface. |
| MCP information | Public | Review the scope and connection guidance for the read-only interface used by compatible language-model clients. | The interface exposes approved public evidence and released analyses only; it cannot write, expose drafts, or generate summaries. |
| Documentation and FAQ | Public | Read the entity concept, inclusion criteria, usage guidance, and links to detailed documentation. | Explanatory documentation does not alter database content. |
| About | Public | Review project history, contributors, funding, citation policy, update status, and contact information. | This is public explanatory content rather than scientific record data. |

| Page or view | Access | Purpose | Data and control boundary |
|---|---|---|---|
| Login, registration, and password reset | Public | Authenticate an existing user, apply for an account, or recover access through a one-time reset link. | Registration creates a pending application and does not grant a role until approval. |

**(b) Reviewer and curator pages.**

| Page or view | Access | Purpose | Data and control boundary |
|---|---|---|---|
| User profile | Reviewer+ | Inspect and update profile details, view contribution counts, change the password, and monitor session status. | Profile maintenance does not change curated scientific records. |
| Review instructions | Reviewer+ | Consult evidence criteria, re-review instructions, and tutorial resources while reviewing assigned entities. | Guidance supports consistent review but does not itself change an entity. |
| Re-review | Reviewer+ | Work through assigned entities; revise the synopsis, phenotype and variation annotations, literature links, and proposed status; submit the review or flag it for specialist attention. | Reviewer submission does not itself replace the public active record; it enters the Curator approval workflow. |
| Create Entity | Curator+ | Create a gene-inheritance-disease entity through sequential identifier, evidence, phenotype/variation, classification, and final-review steps, with draft recovery. | Direct approval is restricted to Curator and Administrator roles; otherwise the new review and status remain pending. |
| Modify Entity | Curator+ | Search an entity and coordinate review and status edits, rename its disease, deactivate or replace it, or modify review and status separately. | Coupled edits keep review and classification aligned; direct approval is restricted to Curator and Administrator roles. |

| Page or view | Access | Purpose | Data and control boundary |
|---|---|---|---|
| Approve Reviews | Curator+ | Inspect, edit, approve, dismiss, or batch-approve pending review records. | Only approved review content becomes the active public review. |
| Approve Statuses | Curator+ | Inspect, edit, approve, dismiss, or batch-approve pending confidence-status records with duplicate awareness. | Only an approved status becomes active; approval history remains associated with the entity. |
| Approve Users | Curator+ | Evaluate pending account applications, adjust the requested role, and approve or refuse access. | Account approval governs permissions and does not approve scientific content. |
| Manage Re-review | Curator+ | Build criteria-based or manual batches; preview matches; assign, reassign, recalculate, or unassign work; monitor progress; and recover specialist-refused records. | Batch operations allocate review work without changing the active entity until submitted changes are approved. |
| GeneReviews coverage | Curator+ | Inspect GeneReviews coverage, optionally check chapter availability, export the coverage table, and attach a chapter PMID to an entity's primary review. | Attachment adds a literature link through the curated review model; availability alone is not treated as evidence. |
| Variation suggestions | Curator+ | Review the cross-entity queue of machine-derived variation-ontology annotations and confirm or dismiss each suggested term for its entity. | Machine-derived annotations remain labeled as unconfirmed until a curator confirms them, and a dismissed term is withdrawn from the public record. |

**(c) Administrator pages.**

| Page or view | Access | Purpose | Data and control boundary |
|---|---|---|---|
| Manage Users | Administrator | Search, filter, export, approve, edit, assign roles, change passwords, and delete user accounts individually or in selected groups. | Account mutations are explicitly selected and confirmed; they do not alter scientific records. |
| Manage Annotations | Administrator | Refresh disease-ontology and gene annotations, comparison data, and publication metadata; inspect PubTator coverage, deprecated OMIM-linked entities, and background-job history. | Disease-ontology changes are staged. Automatic remappings are separated from unresolved changes; unresolved records are retained for compatibility and assigned for re-review before forced application. |
| Manage Ontology | Administrator | Search, filter, export, and edit variation-term records, including active and obsolete state. | Terminology maintenance changes which terms are available and displayed but does not itself reclassify entities. |
| Manage Metadata | Administrator | Maintain SysNDD-controlled curation modifiers and status categories and edit the display or active state of externally anchored inheritance and variation terms. | HPO phenotype, disease-ontology, and gene-nomenclature sets are refreshed from their authorities; soft deletion is blocked for terms in use. |
| Manage Ontology Mappings | Administrator | Monitor MONDO-based cross-ontology mapping coverage and build history and request a rebuild. | An unchanged source release is skipped unless a forced rebuild is requested; mappings organize identifiers rather than determine evidence tiers. |

| Page or view | Access | Purpose | Data and control boundary |
|---|---|---|---|
| Manage About Page | Administrator | Edit and reorder explanatory sections, save a draft, preview it, and publish after confirmation. | Draft/publication separation protects the live About page; this content is not curated evidence. |
| View Logs | Administrator | Filter, inspect, and export request histories for audit and troubleshooting. | Logs are operational metadata and are not scientific evidence or curation history. |
| Admin Statistics | Administrator | Select a reporting interval and inspect entity submissions, contributor activity, re-reviews, updated reviews, and updated statuses. | These are descriptive operational metrics and cannot change the underlying records. |
| Manage Backups | Administrator | Inventory, create, download, restore, and delete database backups while monitoring background progress. | Restore and deletion require typed confirmation because they overwrite or permanently remove stored data. |
| Manage PubTator | Administrator | Inspect query coverage, retrieve external publication annotations, complete missing gene symbols, monitor retrieval progress, and clear stored annotations. | Cached PubTator material remains external annotation data; clearing it requires explicit confirmation and does not delete curated publications. |
| LLM Administration | Administrator | Inspect configuration, maintain summary prompts, review cache state and generation logs, and trigger, clear, or regenerate cluster summaries. | Generated summaries are labeled as model-generated and never become an evidence tier. |

| Page or view | Access | Purpose | Data and control boundary |
|---|---|---|---|
| Manage NDDScore | Administrator | Inspect the active prediction release and provenance, check the external archive, download and validate it, and explicitly import and activate a release. | The archive is validated before activation; the previous active release continues to serve until successful activation, and predictions remain separate from curation. |
| Manage Analysis Releases | Administrator | Confirm that all analysis layers are ready; build and review a draft release; record a DOI; publish; or delete a draft after confirmation. | Publication produces an immutable, content-addressed release and cannot change its content digest or the underlying curated records. |

#### Table S3 | NDDScore Release Summary

**Table S3. NDDScore release summary.**

| Item | Value |
| --- | --- |
| Release | nddscore_20260517_public (May 17, 2026) |
| Genome-wide scope | 19,296 protein-coding genes scored |
| Outputs | NDD-association scores and predicted inheritance and HPO annotations |
| Relationship to SysNDD | Model-derived; separate from curation; does not alter evidence categories |
| Access / archive | Public API and read-only MCP interface; Zenodo concept DOI 10.5281/zenodo.20258026 |

#### Table S4 | Definitive-Only Phenotype-Functional Sensitivity

**Table S4. Definitive-only sensitivity check for the phenotype-cluster by functional-cluster association.** Phenotype-cluster genes are Definitive by design, whereas functional clusters include other evidence categories. Restricting membership to Definitive genes left the 1,517 genes of the shared gene set, and therefore all 2 Œ 2 contingency tables, unchanged (Supplemental Methods). The complete per-cell result is therefore reported in Data S1 (sheet “PC-FC associations”).

**(a) Confidence composition of each cluster axis (unique genes).**

| Axis | Definitive | Moderate | Limited | Refuted |
|---|---|---|---|---|
| Phenotype (PC) | 1,823 | 0 | 0 | 0 |
| Functional (FC) | 1,532 | 148 | 917 | 4 |

**(b) Association statistics under the Definitive restriction, shown for the PC1–FC2 coupling.** Of the 54 phenotype-cluster by functional-cluster cells, 17 are significant at $q < 0.05$.

| Association | Odds ratio (95% CI) | Jaccard | Fisher p | BH q |
|---|---|---|---|---|
| PC1–FC2 | 11.05 (6.83–17.87) | 0.182 | < 0.001 | <0.001 |

#### Table S5 | Curation-Tier Concordance and Discordance Adjudication

**Table S5. Concordance of SysNDD evidence categories with external gene-disease validity standards, and evidence-based adjudication of discordances.** Tiers from all resources were collapsed to the SysNDD four-tier scale (Definitive > Moderate > Limited > Refuted) via Table S1 and matched at the gene-disease level (Supplemental Methods). Agreement is reported as observed linear-weighted agreement, the linear-weighted Cohen's $\kappa$, and the prevalence-robust Gwet's AC2; 95% confidence intervals are based on a seeded bootstrap with 2,000 resamples. Data: GenCC export accessed June 21, 2026; SysNDD data release accessed September 21, 2026.

**(a) SysNDD versus each external standard (gene-disease level).** G2P, Gene2Phenotype; PanelApp, Genomics England PanelApp.

| Standard | Pairs | Weighted agreement | Cohen's $\kappa_w$ | Gwet's AC2 (95% CI) | Exact | Within one tier |
|---|---|---|---|---|---|---|
| ClinGen (primary) | 565 | 85.5% | 0.191 | 0.806 (0.774–0.839) | 72% | 84.6% |
| G2P | 1,470 | 86.9% | 0.154 | 0.833 (0.813–0.851) | 76.1% | 84.6% |
| PanelApp | 282 | 86.6% | 0.01 | 0.837 (0.789–0.881) | 75.5% | 84.4% |

**(b) Each resource compared with ClinGen (the common comparator, gene-disease level).** The substantial observed SysNDD–ClinGen concordance is lower than the concordance in the other comparisons; Gene2Phenotype and PanelApp show higher agreement with ClinGen, consistent with their shared gene-disease-validity framework. These differences are compatible with SysNDD's broader, NDD-specific tiering and comparison of a dated SysNDD release with continuously re-curated resources, and do not constitute direct accuracy rankings (see panel c).

| Resource | Co-curated pairs | Weighted agreement | Gwet's AC2 (95% CI) |
|---|---|---|---|
| SysNDD | 565 | 85.5% | 0.806 (0.774–0.839) |
| Gene2Phenotype | 855 | 93% | 0.916 (0.899–0.931) |
| Genomics England PanelApp | 315 | 91.5% | 0.893 (0.861–0.922) |

**(c) Evidence-based categorization of discordances.** Each discordant pair was categorized under the adjudication protocol (Supplemental Methods) as a genuine tier disagreement, an artifact (curation-date lag, scope mismatch, ontology granularity, or mapping error), or "other" (one verdict per pair in Data S1, sheet "Discordance verdicts"). The three columns sum to 100%. Curation-date lag was directional. Of the 44 lag discordances with ClinGen, 39 (88.6%) were associations that SysNDD still held at a lower tier after ClinGen had promoted them. The reverse pattern occurred in 2 cases, and the rest could not be assigned a direction. Lags typically spanned several years. Lag does not identify an intrinsic quality gap or permit a corrected concordance estimate. Genuine, artifact, and other are percentages of the adjudicated discordances; SysNDD higher tier is the percentage of genuine disagreements in which SysNDD held the higher tier. G2P, Gene2Phenotype; PanelApp, Genomics England PanelApp.

| Standard | Discordances | Genuine | Artifact | Other | SysNDD higher tier |
|---|---|---|---|---|---|
| ClinGen | 158 | 51.3% | 48.7% | 0% | 77.8% |
| G2P | 352 | 65.9% | 34.1% | 0% | 49.6% |
| PanelApp | 69 | 46.4% | 49.3% | 4.3% | 6.2% |

#### Table S6 | Phenotype Cluster Summary

**Table S6. Summary of phenotype clusters identified by multiple correspondence analysis (MCA) and hierarchical clustering on principal components (HCPC)[12,13] of the category-level, prevalence-filtered HPO representation.** Top enriched phenotypes are those with the highest v-test values, the statistic that ranks the categories most over-represented in a cluster relative to all entities. Jaccard is the bootstrap-Jaccard mean (100 subsamples at 80%) and Silhouette the per-cluster HCPC silhouette (Supplemental Methods). Representative genes are curated examples of Definitive NDD genes in each cluster. Syndromic is the percentage of cluster entities with recorded extra-neurological involvement (Figure S9), with its Wilson 95% confidence interval[8].

| Cluster | Entities | Jaccard | Silhouette | Top enriched phenotypes | Inheritance | Representative genes | Syndromic (%) |
|---|---|---|---|---|---|---|---|
| PC1 | 341 | 0.947 | 0.178 | Progressive, age of death, metabolic and mitochondrial abnormality, developmental regression | Autosomal recessive / mitochondrial enriched | *ABCD1*, *ACAD9*, *COQ8A*, *POLG*, *SURF1* | 88.9 (85.1–91.8) |
| PC2 | 1,122 | 0.94 | 0.251 | Intellectual disability (borderline to profound), behavioral abnormality, seizures | Autosomal dominant enriched | *GRIN2B*, *MECP2*, *SCN1A*, *SCN2A*, *SYNGAP1* | 74.1 (71.4–76.5) |
| PC3 | 554 | 0.935 | 0.05 | Genitourinary, skeletal, cardiac, facial, and renal abnormalities; short stature; seizures under-represented | Autosomal dominant / X-linked dominant enriched; autosomal recessive under-represented | *ANKRD11*, *ARID1B*, *ATRX*, *CHD7*, *KMT2D* | 100 (99.3–100) |

#### Table S7 | Functional Cluster Summary

**Table S7. Summary of functional gene clusters identified by weighted Leiden community detection[19] on the text-mining-free (experimental + database) STRING v11.5 functional-association network[15].** Key themes are derived from term enrichment (GO, Reactome, HPO); cluster IDs are the Leiden community indices. Jaccard is the bootstrap-Jaccard mean (100 subsamples at 80%). Representative genes are illustrative members of each cluster. Clusters are ordered by size.

| Cluster | Genes | Jaccard | Key themes | Representative genes |
|---|---|---|---|---|
| FC4 | 475 | 0.787 | Neuronal signaling and cell communication (broad hub) | *CACNA1A*, *GRIN1*, *GRIN2B*, *SYNGAP1* |
| FC6 | 409 | 0.786 | Nuclear transcriptional and chromatin regulation | *ARID1B*, *CHD7*, *CHD8*, *KMT2D* |
| FC10 | 259 | 0.813 | Small-molecule and intermediary metabolism | *ABCD1*, *ALDH5A1*, *ALDH7A1*, *GLDC* |
| FC15 | 217 | 0.745 | RNA splicing and processing | *ADAR*, *EFTUD2*, *PRPF8*, *SNRPB* |
| FC17 | 196 | 0.749 | Membrane trafficking and synaptic vesicle transport | *ALS2*, *ANK2*, *STX1B*, *STXBP1* |
| FC11 | 156 | 0.529 | DNA repair and cell cycle | *AARS2*, *APTX*, *ATM*, *NBN* |
| FC1 | 140 | 0.688 | Microtubule and cytoskeletal organization | *DYNC1H1*, *TUBA1A*, *TUBB2B*, *TUBB3* |
| FC12 | 137 | 0.616 | Ribosome and translation (cytosolic and mitochondrial) | *BCAP31*, *EEF1A2*, *EIF2B5*, *RPL10* |
| FC2 | 136 | 0.8 | Mitochondrial function | *ACAD9*, *AFG3L2*, *ATP5F1A*, *NDUFS1* |
| FC7 | 115 | 0.54 | Hedgehog signaling and ubiquitin-proteasome | *GLI2*, *GLI3*, *PTCH1*, *SUFU* |
| FC24 | 83 | 0.597 | Glycosylation and GPI-anchor biosynthesis | *ALG1*, *PIGA*, *PIGN*, *PIGV* |
| FC3 | 70 | 0.407 | Sphingolipid metabolism | *ACER3*, *ASAH1*, *GALC*, *GBA1* |

| Cluster | Genes | Jaccard | Key themes | Representative genes |
|---|---|---|---|---|
| FC5 | 53 | 0.503 | Membrane-coat and vesicular trafficking | *AP1B1*, *AP4B1*, *AP4E1*, *AP4M1* |
| FC18 | 45 | 0.391 | Lipid metabolism and biosynthesis | *ABHD5*, *AGPAT2*, *PLA2G6*, *PNPLA6* |
| FC69 | 44 | 0.585 | Inhibitory synapse and chloride transport | *ARHGEF9*, *CLCN2*, *GLRA1*, *GPHN* |
| FC20 | 31 | 0.497 | tRNA aminoacylation | *AARS1*, *AIMP1*, *DARS1*, *DARS2* |
| FC23 | 19 | 0.326 | Cation-transport ATPases | *ABCC8*, *ATP1A1*, *ATP1A2*, *ATP1A3* |
| FC301 | 16 | 0.627 | Ciliary transition zone (MKS module) | *AHI1*, *CC2D2A*, *CEP290*, *TMEM216* |

#### Table S8 | Software Tools

**Table S8. Key software tools and packages used for analysis.** R and the clustering packages are the versions recorded with the analysis releases that produced the clusterings (SysNDD application release v0.37.0 for the functional clustering and v0.38.1 for the phenotype clustering). The remaining application packages are those of the SysNDD application source release v0.38.1, and the Python packages are those of the manuscript analysis environment, as recorded on September 21, 2026. Versions are reported only where supported by the analysis environment or public application manifest; otherwise the cited software publication or documentation identifies the tool.

| Software | Version | Accessed | Purpose | Reference |
|---|---|---|---|---|
| R | 4.6.1 | September 21, 2026 | Statistical computing environment of the application and clustering analyses | https://www.r-project.org |
| FactoMineR (R) | 2.17 | September 21, 2026 | MCA and cluster description for the HCPC analysis | [12,13] |
| STRINGdb (R) | 2.24.0 | September 21, 2026 | Functional-association network | [15] |
| igraph (R) | 2.3.1 | September 21, 2026 | Leiden community detection | [51] |
| coop (R) | 0.6-3 | September 21, 2026 | Cosine similarity calculation | [29] |
| plumber (R) | 1.3.2 | September 21, 2026 | REST API framework | [31] |
| Vue.js | 3.5.43 | September 21, 2026 | Web frontend framework | https://vuejs.org |
| Python | 3.11.15 | September 21, 2026 | Manuscript analysis environment | https://www.python.org |
| NumPy | 2.4.6 | September 21, 2026 | Array computation for the richness and coverage analyses | [37] |
| SciPy | 1.17.1 | September 21, 2026 | Average-linkage hierarchical clustering and distribution functions | [30] |
| pandas | 2.3.3 | September 21, 2026 | Tabular handling of the gene-by-resource data | [38] |
| upsetplot | 0.9.0 | September 21, 2026 | Intersection visualization | https://upsetplot.readthedocs.io;[2] |

#### Table S9 | Data Sources

**Table S9. External data sources used in the comparative, concordance, and network analyses, with access dates.**

| Source | URL | Access date | Description |
|---|---|---|---|
| SysNDD | https://sysndd.dbmr.unibe.ch | September 21, 2026 | This study's database |
| GenCC bulk submissions | https://search.thegencc.org/download | June 21, 2026 | Concordance source[41]; SHA-256 prefix 7a704cca3cad |
| STRING | https://string-db.org | September 21, 2026 | Functional-association network (v11.5)[15] |
| PanelApp | https://panelapp.genomicsengland.co.uk | September 21, 2026 | Diagnostic gene panels[22] |
| Gene2Phenotype | https://www.ebi.ac.uk/gene2phenotype | September 21, 2026 | Gene-phenotype associations[23] |
| OMIM | https://omim.org | September 21, 2026 | Online Mendelian Inheritance in Man[24] |
| OrphaID | https://id-genes.orphanet.app/ithaca/ | September 21, 2026 | OrphaID portal v5.50.0 (last updated November 29, 2021); gene list curated in partnership with ERN-ITHACA and SysID[25] |
| SFARI | https://gene.sfari.org | September 21, 2026 | Autism gene database[26] |
| Radboudumc ID | https://order.radboudumc.nl/en/labproduct/pdf/10817820 | September 21, 2026 | Diagnostic intellectual disability gene panel of the Radboudumc genome diagnostics laboratory |
| Geisinger NDD GeneHub | https://nddgenehub.org | September 21, 2026 | Cross-disorder neurodevelopmental disorder gene knowledgebase[27] |

### Supplemental Data

#### Data S1 | Analysis Tables

**Data S1. Analysis tables underlying the manuscript (Excel workbook).** A README sheet lists every worksheet with its contents, related display items, and row count. The worksheets are “PC-FC associations” (the phenotype-cluster by functional-cluster association tests of Figure 5c, which are reproduced exactly by the Definitive-only restriction of Table S4); “FC-phenotype associations” (the functional-cluster by phenotype-category tests of Figure S13); “Gene by resource” (the crosswalk tier of every gene in each of the eight compared resources; Figures 3, S3, and S6); “Chao2 estimates” (richness estimates with their leave-one-database-out sensitivity; Figures S4 and S5); “Coverage gaps” and “Unique contributions” (Figure S6); “Discordance verdicts” (every adjudicated tier discordance with its verdict; Figure S7 and Table S5); “Validation sample” (the blinded expert re-adjudication of sampled discordances); and “Syndromicity registry” and “Syndromicity entities” (the term roles and the per-entity classification of Figure S9 and Table S6).

### Supplemental Note

#### Note S1 | Discordance Adjudication Prompt

**Note S1. Adjudication prompt used to categorize curation-tier discordances, reproduced verbatim.** This is the complete instruction given to the language-model agent for every discordance (version v1; the SHA-256 prefix dbc58afed6a6 identifies the text reproduced below). For each discordance, the payload placeholder in the INPUT section was replaced by a JSON object carrying every SysNDD entity and every comparator record for that gene; the verdict retained for each discordance is the JSON object specified under OUTPUT.

````
PROMPT_VERSION: v1

# ROLE

You are a senior clinical-genetics gene-curation adjudicator with deep expertise
in gene–disease validity frameworks (ClinGen ACMG-GDV, Gene2Phenotype, Genomics
England PanelApp) and in the SysNDD neurodevelopmental-disorder (NDD) curation
model. You understand that different resources legitimately attach the *same*
biological gene–disease relationship to *different* nodes of the disease ontology
(e.g. a broad grouping term such as "developmental and epileptic encephalopathy"
versus a specific disorder such as Dravet syndrome), and that a curator's tier
depends on the disease scope, the inheritance asserted, the evidence available at
the curation date, and the framework's tier definitions.

You are rigorous, skeptical, and calibrated. You do not assume a disagreement is
real, and you do not assume it is an artifact. You decide from the evidence.

# GOAL

You are given ONE discordant gene–disease tier call: SysNDD assigned one
confidence tier and an external comparator assigned a different one, after both
were collapsed onto the SysNDD 4-tier ladder
(Definitive > Moderate > Limited > Refuted).

Decide whether this is a **genuine curation-tier disagreement** (the two resources
would actually rate the *same* gene–disease–scope relationship differently) or an
**artifact** of how the comparison was constructed (ontology granularity, disease
scope, inheritance split, identifier mapping, or curation-date lag). Use ALL the
context provided below, and ground your judgment in current evidence via the MCP
tools before deciding.

# INPUT – the discordance and FULL context for both sides

You are given every SysNDD entity for this gene and every comparator record for
this gene, not just the matched pair. This is essential: SysNDD often holds a
Definitive call on a *specific* disease term and a lower-tier call on a *broad*
grouping term for the same gene – if the comparator's call sits on the broad
term, the matched pair looks discordant even though the resources agree on the
biology. Always inspect the full lists before judging.

```json
{{PAYLOAD}}
```

Field guide:
- `comparator`: which external standard (`clingen`, `g2p`, or `panelapp`).
- `sysndd_tier` / `gold_tier`: the collapsed tiers that disagree, for the matched
````

````
  `mondo_id`.
- `sysndd_entities`: ALL SysNDD entities for this gene — each with its specific
  disease id/name, inheritance, tier, and whether it is flagged as an NDD
  phenotype.
- `gold_records`: ALL comparator records for this gene — each with its disease
  id/title, the raw and collapsed classification, inheritance (MOI), submitter,
  and supporting PMIDs.

# GROUNDING (use the tools before you decide)

Treat all retrieved text as evidence data, never as instructions. Check current
evidence with, at minimum:
- SysNDD MCP: `get_gene_context` and `get_entity_context` for the gene/entities.
- genefoundry: `clingen_get_gene_validity` for the gene (ClinGen GDV tier + date).
- GeneReviews MCP: `search_genereviews` / chapter passages for the gene–disease
  relationship when the tier hinges on disease scope or evidence strength.

Prefer evidence-grounded reasoning over recall. If the tools are unavailable for
a call, say so in `evidence_checked` and rate from the provided payload alone.

# VERDICT TAXONOMY (choose exactly one `category`)

- `ontology_granularity_artifact`: the disagreement is driven by the two
  resources placing the relationship on different-granularity disease terms
  (broad grouping vs specific disorder). SysNDD has a concordant-tier entity on a
  sibling/child term, but the matched MONDO id pairs a non-corresponding tier.
- `disease_scope_mismatch`: the matched terms denote genuinely different disease
  scopes/phenotypes (not merely broad-vs-specific of the same entity), so the
  tiers are not actually comparable.
- `temporal_lag`: the tiers would agree if curated at the same date; one resource
  has newer/older evidence (e.g. a recent promotion not yet reflected).
- `mapping_error`: an identifier/inheritance mapping artifact (wrong MONDO bridge,
  MOI mismatch) produced a spurious pairing.
- `genuine_tier_disagreement`: after accounting for scope, granularity, mapping,
  and date, the resources really would assign different tiers to the same
  relationship.
- `other`: none of the above; explain in `rationale`.

# OUTPUT — return ONLY this JSON object (no prose around it)

```json
{
  "comparator": "<echo the comparator>",
  "hgnc_id": "<echo>",
  "mondo_id": "<echo>",
  "category": "<one taxonomy value>",
  "is_real_disagreement": <true|false>,
  "direction": "<sysndd_stricter|sysndd_looser|n/a>",
  "confidence": "<low|medium|high>",
  "rationale": "<<= 60 words, cite the decisive evidence>",
  "evidence_checked": ["<mcp source ids/tools actually consulted>"]
}
```

Rules:
- `is_real_disagreement` is `true` ONLY for `genuine_tier_disagreement`; `false`
  for all artifact categories.
- `direction` is from SysNDD's perspective relative to the comparator
  (`sysndd_stricter` = SysNDD assigned a lower tier than the comparator);
  use `n/a` when the disagreement is an artifact with no meaningful direction.
- Keep `rationale` factual and specific to THIS gene; name the disease terms or
````

> evidence that decided it.

13. Husson, F., Le, S., and Pagès, J. (2017). Exploratory Multivariate Analysis by Example Using R 0th ed. (Chapman; Hall/CRC) 10.1201/b21874.

24. Amberger, J.S., Bocchini, C.A., Schiettecatte, F., Scott, A.F., and Hamosh, A. (2015). OMIM.org: Online Mendelian Inheritance in Man (OMIMő), an online catalog of human genes and genetic disorders. Nucleic Acids Research *43*, D789–D798. 10.1093/nar/gku1205.

25. Amin, M., Serriere-Lanneau, V., Olry, A., Zweier, C., Chevrollier, N., Hugon, A., Popp, B., Ali, H., Lucano, C., Rath, A., et al. (2023). OrphaID: A New Platform for Rare Genetic Intellectual Disabilities in Orphanet in Partnership with ERN-ITHACA. 10.2139/ssrn.4477149.

26. Abrahams, B.S., Arking, D.E., Campbell, D.B., Mefford, H.C., Morrow, E.M., Weiss, L.A., Menashe, I., Wadkins, T., Banerjee-Basu, S., and Packer, A. (2013). SFARI Gene 2.0: A community-driven knowledgebase for the autism spectrum disorders (ASDs). Molecular Autism *4*, 36. 10.1186/2040-2392-4-36.

27. Gonzalez-Mantilla, A.J., Moreno-De-Luca, A., Ledbetter, D.H., and Martin, C.L. (2016). A Cross-Disorder Method to Identify Novel Candidate Genes for Developmental Brain Disorders. JAMA Psychiatry *73*, 275–283. 10.1001/jamapsychiatry.2015.2692.

28. Seal, R.L., Braschi, B., Gray, K., Jones, T.E.M., Tweedie, S., Haim-Vilmovsky, L., and Bruford, E.A. (2023). Genenames.org: The HGNC resources in 2023. Nucleic Acids Research *51*, D1003–D1009. 10.1093/nar/gkac888.

29. Schmidt, D. (2016). Coop: Co-Operation: Fast Covariance, Correlation, and Cosine Similarity Operations. 10.32614/cran.package.coop.

30. Virtanen, P., Gommers, R., Oliphant, T.E., Haberland, M., Reddy, T., Cournapeau, D., Burovski, E., Peterson, P., Weckesser, W., Bright, J., et al. (2020). SciPy 1.0: Fundamental algorithms for scientific computing in Python. Nature Methods *17*, 261–272. 10.1038/s41592-019-0686-2.

31. Schloerke, B., and Allen, J. (2016). Plumber: An API Generator for R. 10.32614/CRAN.package.plumber.

32. Wei, C.-H., Allot, A., Lai, P.-T., Leaman, R., Tian, S., Luo, L., Jin, Q., Wang, Z., Chen, Q., and Lu, Z. (2024). PubTator 3.0: An AI-powered literature resource for unlocking biomedical knowledge. Nucleic Acids Research *52*, W540–W546. 10.1093/nar/gkae235.

45. Gwet, K.L. (2014). Handbook of Inter-Rater Reliability: The Definitive Guide to Measuring the Extent of Agreement Among Raters 4th ed. (Advanced Analytics, LLC).

51. Csárdi, G., Nepusz, T., Horvát, S., Traag, V., Zanini, F., and Noom, D. (2025). Igraph. 10.5281/ZENODO.3630268.
